# A Fkh1/2 binding site array in the *WHI5* promoter drives sub-scaling transcription

**DOI:** 10.1101/2025.10.10.681508

**Authors:** Jacob Kim, Shicong Xie, Lucas Fuentes Valenzuela, Jordan Xiao, Mike C. Lanz, Xin Gao, Matthew Swaffer, Corbin E. Mitchell, Seth M. Rubin, Kurt Schmoller, Jan M. Skotheim

## Abstract

Cells typically regulate their size within a relatively tight range by coupling growth to the cell division cycle using a dedicated set of molecular mechanisms. In budding yeast, cells are born with a similar amount of the G1/S inhibitor protein Whi5 that is then diluted by growth throughout G1. As cells grow, Whi5 concentration decreases and cells become more likely to enter the cell cycle. Cells are born in G1 with similar amounts of Whi5 because of the size-independent (sub-scaling) expression of *WHI5* mRNA during S/G2/M phases and the equal partitioning of Whi5 protein at division. While the latter is known to be achieved by association with chromatin before anaphase, the mechanism for the former is poorly understood. Through systematic mutations of the *WHI5* promoter, we discovered that *WHI5*’s core promoter region located -126 to -75 base pairs upstream of the start codon is responsible for sub-scaling expression. This sequence contains a repeating array of binding sites for the transcription factors Fkh1 and Fkh2. Mutation of any of these sites, deletion of either *FKH1* or *FKH2*, or preventing Fkh1 or Fkh2 dimerization weakens the sub-scaling of *WHI5* transcription. Taken together with structural predictions and a mathematical model of cooperative Fkh-DNA binding, we conclude that *WHI5*’s sub-scaling transcription is regulated by a Fkh1/2 heteropolymer binding an array of sites in its core promoter.

## INTRODUCTION

Cell size is fundamental to cellular physiology as it is tightly linked to biosynthetic capacity, nutrient uptake, and organelle scaling and function^1–4^. Consistent with the idea that precise size control is essential for normal cellular function, most cells maintain a characteristically tight size range. Deviations from this range can impair cellular function. For example, if cells grow excessively large, many cytoplasmic components become diluted and cells both grow inefficiently and become senescent^5–8^. Thus, understanding how cells sense and regulate their size is central to cell biology.

Proliferating cells typically control their size by linking cell growth to the cell division cycle. In the organisms where cell size control has been most actively studied, it has become increasingly clear that there are specific signaling pathways devoted to sensing cell size and using this information to regulate cell division^4,9^. One way cells sense their size is to link the concentration of specific regulators of cell division to cell size^10^. In the fission yeast *S. pombe*, the concentration of the Cdc25 phosphatase that triggers mitosis increases with cell size^11^, while in the budding yeast *S. cerevisiae*, the G1/S transition inhibitor Whi5 is diluted by cell growth in the G1 phase of the cell cycle^12–14^. This type of dilution of cell cycle inhibitors in the G1 phase of the cell cycle has also been found to occur in mammalian cells through the dilution of the retinoblastoma protein Rb1^15–17^, in *A. thaliana* plants through the dilution of the F-box protein KRP4^18^, and in *C. reinhardtii* green algae through the dilution of the mRNA binding protein TNY1^19^. In contrast, other cell cycle regulators examined in these organisms remained at approximately constant concentration as cells grew larger. Thus, a common feature of cell size regulation across eukaryotes is the change in concentration of specific cell cycle regulators with increasing cell size to promote division^4^.

The identification of size-dependent concentration changes as drivers of cell division raised the question as to what are the underlying mechanisms that drive these changes. In human cells, the concentration of Rb in G1 is regulated by the degradation of its hypo-phosphorylated isoforms^17^. This degradation stops once Rb is hyper-phosphorylated at the G1/S transition, which allows it to reaccumulate during the S/G2/M phases to prepare to regulate the subsequent cell cycle. In contrast to Rb, accumulation of Cdc25 in fission yeast and the dilution of Whi5 in budding yeast reflect their transcriptional regulation^11,13^ (**Figure 1A**). In the case of budding yeast, Whi5 is mostly synthesized in the S/G2/M phases of the cell cycle due to its cell cycle-dependent transcriptional regulation^12,20^. Then, the stable Whi5 protein is diluted by cell growth during G1. The synthesis of Whi5 is at a rate approximately independent of cell size so that big and small cells make approximately the same amount of protein in S/G2/M phases of the cell cycle. This largely size-independent synthesis of Whi5 protein arises from the largely size-independent transcription of *WHI5* mRNA, whose concentration sub-scales and is lower in larger cells^13^. Whi5 binds to chromatin during mitosis to be equally partitioned into the larger mother cell and the smaller daughter cell^13^. Thus, sub-scaling expression and chromatin-based partitioning of Whi5 protein ensures that all cells inherit similar amounts at division, preserving the inverse correlation between Whi5 concentration and cell size. Disrupting both these mechanisms reduces the ability of a cell to control its size^13^.

**Figure 1:**
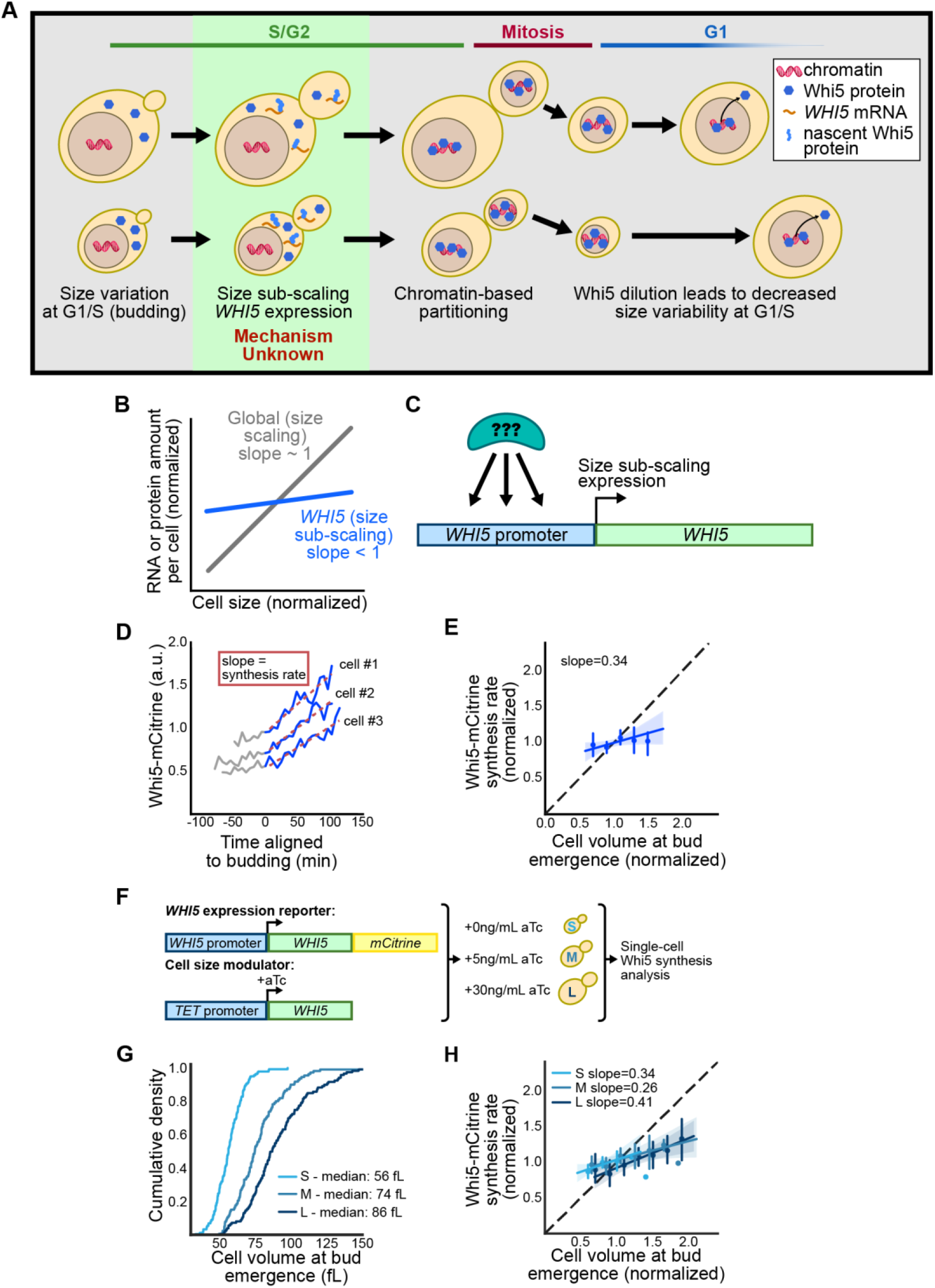
Whi5 synthesis sub-scales with cell size without feedback. (A) Schematic diagram illustrating how the G1 size control and sub-scaling of Whi5 protein expression arises from the sub-scaling transcription in S/G2/M phases followed by equal chromatin-based partitioning into the larger mother and smaller daughter cells at division. (B) Sub-scaling proteins are defined as those whose amounts increase less with cell size than the total amount of protein, which scales in proportion to cell size. (C) The *WHI5* promoter is responsible for its sub-scaling expression through an unknown mechanism. (D) Characteristic single cell traces of the amount of Whi5-mCitrine fluorescent protein expressed from the endogenous *WHI5* promoter through a cell cycle. A linear fit to the protein amount traces in the budded period of the cell cycle (S/G2/M) was used to determine the protein synthesis rate for each individual cell. (E) Whi5-mCitrine synthesis rates normalized to the average rate and plotted as a function of cell size at bud emergence, which was normalized to the average. The linear fit with a slope < 1 indicates sub-scaling expression. The line denotes a linear fit and error bars and shaded area indicate 95% confidence intervals. (n=140) (F) Schematic illustrating the aTc-regulated synthetic promoter used to express exogenously controlled *WHI5* used to modulate average cell size. (G) Cumulative distributions for the cell size at budding for the small (n=140), medium (n=111), and large (n=120) cell populations that are exposed to 0, 5, and 30ng/mL aTc, respectively. (H) Normalized Whi5-mCitrine synthesis rate for the small (n=140), medium (n=111), and large (n=120) cells as in (E). A multivariate regression did not identify a significant effect of exogenous Whi5 on the synthesis rate of the endogenous Whi5-mCitrine (p>0.05 all comparisons). The lines denote linear fits and error bars and shaded areas indicate 95% confidence intervals.

That Whi5’s concentration during G1 is inversely related to cell size reflects both its chromatin based partitioning and sub-scaling transcription^13^. While the binding mechanisms through which Whi5 associates with the DNA-bound SBF transcription factor to enable its partitioning have been identified^21^, we knew little of the transcriptional mechanism that ensures the synthesis of Whi5 occurs at a similar rate in large and small cells. Here, show that the core promoter of *WHI5*, located -126 to -76 base pairs upstream of the start codon, is responsible for its sub-scaling transcription. This core promoter contains an array of binding sites for the Fkh1 and Fkh2 transcription factors. Removal of any of these sites, deletion of either *FKH1* or *FKH2*, or introducing dimerization-deficient mutations to either *FKH1* or *FKH2* reduces the size-independence of *WHI5* transcription. Taken together with structural and mathematical models, our data suggest that the binding of a Fkh1/2 heteropolymer to the Fkh1/2 binding site array is responsible for the sub-scaling of *WHI5* transcription.

## RESULTS

### Exogenously controlled Whi5 does not impact the expression of endogenous Whi5

The synthesis of similar amounts of Whi5 in large and small cells arises from the fact that they spend similar amounts of time in the S/G2/M phases of the cell cycle and synthesize Whi5 protein at a similar rate during this time^12,22^. This similar Whi5 synthesis rate was shown to arise at the level of transcription as both large and small S/G2/M phase cells had similar numbers of *WHI5* mRNA^13^ (**Figure 1B**). In this way, the expression of *WHI5* sub-scales with cell size, meaning that its concentration is lower in larger cells^14^. In contrast, the expression of most yeast mRNAs is closer to scaling in proportion to the size of the cell so that their concentrations remain roughly the same as cells grow larger^23^. This more unique sub-scaling relationship between *WHI5* expression and cell size is due to its promoter, which was found to be both necessary and sufficient for sub-scaling synthesis. Namely, the expression of Whi5-mCitrine from an *ACT1* promoter increased in proportion to cell size, while the expression of mCitrine from a 1kbp *WHI5* promoter did not^13^.

To identify the mechanism through which the *WHI5* promoter drives size-independent expression, we decided to make a series of mutations to the promoter and then measure their effects on the size-dependence of the expression. We previously assessed the size-dependence of gene expression by measuring the rate of synthesis (accumulation) of a stable fluorescent protein in single cells (see **methods**). Since different cells entered S phase at different sizes, we could estimate the correlation of cell size at budding and the rate of synthesis of the stable fluorescent protein^12^ (**Figures 1D-E, S1A**). However, accurately estimating this size-correlation and assessing the differences between strains with mutations in their *WHI5* promoters is difficult because of the limited natural variation in cell sizes at the onset of S phase.

Since we anticipated measuring the size-correlation of expression in many promoter mutants we sought a more efficient way of measuring size-scaling by expanding on the natural variation of cell size at the onset of S phase. To do this, we expressed additional Whi5 protein from a synthetic *TET* promoter that is conditionally activated by anhydrotetracycline (aTc) in a dose-dependent manner^24^ (**Figures 1F-G, S1B-C**). Exposing cells to 30ng/mL aTc generated a population of larger than normal cells in which we could monitor the expression of endogenous *WHI5pr-WHI5-mCitrine*. Examining the rate of accumulation of Whi5-mCitrine in both populations revealed a similar correlation with cell size indicating that the populations could be pooled together to estimate the size-correlation using fewer cells (**Figures 1H, S1D-E**). This can be seen in a simple linear model predicting the rate of Whi5-mCitrine accumulation as a function of cell size at budding and aTc concentration, where the size at budding is highly statistically significant (p<10^-3^) but the aTc concentration is not (p=0.82 for 5ng/mL aTc, p=0.06 for 30ng/mL aTc). This result supports the view that there is little feedback from Whi5 onto its own expression and is consistent with earlier experiments showing that the rate of Whi5 protein expression in S/G2/M is proportional to the number of copies of the *WHI5* gene^12^. These data also support our use of the exogenously controlled *TET* promoter to express unlabeled Whi5 to expand the dynamic range of cell sizes beyond the natural range typical of wild-type cells.

### The core promoter of *WHI5* is responsible for its sub-scaling gene expression

After establishing that a *TETpr-WHI5* allele can be used to alter cell size without affecting the expression of an endogenous *WHI5pr-WHI5-mCitrine* allele, we decided to use this strain to examine the effect of different *WHI5* promoter mutations on the sub-scaling of its expression. We exposed cells to either no aTc or 30ng/mL aTc to generate populations of mean size 46fL and 69fL, respectively (**Figures 2A and S2A**). In addition to increasing cell size, the exogenous Whi5 led to a slight increase in the fraction of cells in G1 from 34% to 39% (**Figure S2B**). That mutations increasing cell size by inhibiting the G1/S transition only have a modest effect on the fraction of cells in G1 has consistently been observed^25–27^. This fact led us to test if the sub-scaling of *WHI5* mRNA expression we observed previously in S/G2/M phase cells^13^ could be seen in the population as a whole. To do this, we measured relative mRNA concentrations from cells at no aTc and at 30ng/mL aTc using qPCR (see **methods**). We used primers targeting the *mCitrine* sequence to estimate the endogenous *WHI5-mCitrine* concentration and primers targeting the *ACT1* gene whose corresponding mRNA concentration does not change with increasing cell size or cell cycle phase^13,23^. Indeed, this revealed that a 545bp *WHI5pr-WHI5-mCitrine* mRNA sub-scaled with cell size, which is consistent with previous RNA-seq data measuring endogenous *WHI5* mRNA (**Figures 2B and S2C**). The sub-scaling with cell size was likely mainly due to effects on transcription because a similar sub-scaling can be seen in the occupancy of RNA Polymerase II on the *WHI5* gene body^28^ (**Figure 2C**). That the size slopes for RNA Polymerase II occupancy are lower than the size slopes for mRNA may be due to the general increase in mRNA stability seen in larger cells^28^ (**Figure S2D**).

**Figure 2:**
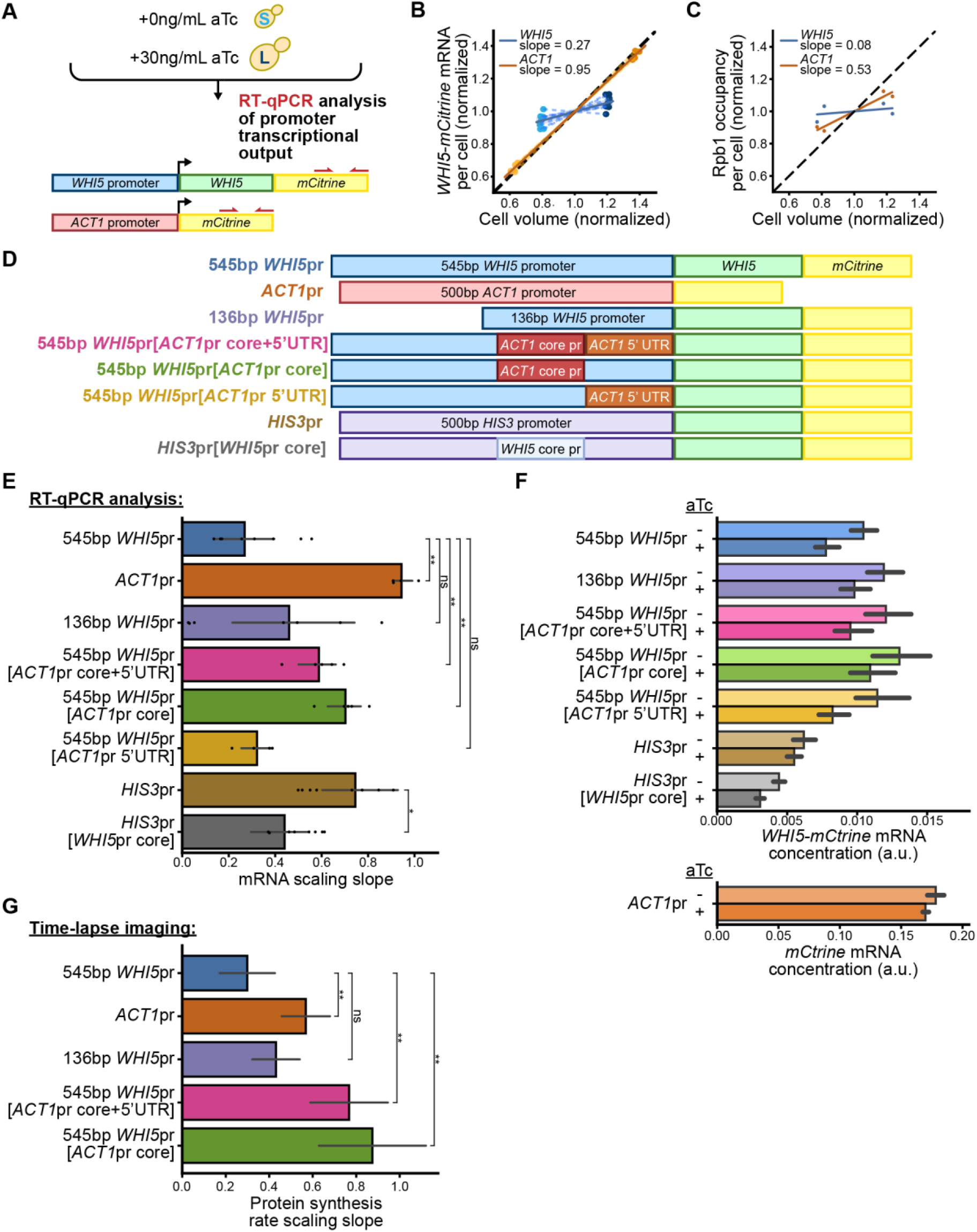
The 51bp *WHI5* core promoter regulates sub-scaling expression. (A) Experiment schematic for RT-qPCR analysis. Red arrows denote the primer pair targeting *mCitrine*, which is fused to the endogenous *WHI5* allele. 0 and 30ng/mL aTc concentrations are used to generate small and large populations of cells, respectively. (B) Scaling slope is defined by the relationship between the relative amount of *WHI5-mCitrine* mRNA per cell measured using RT-qPCR and cell size. The black dashed line indicates perfect size scaling that maintains constant concentration as cells grow larger. Blue dashed lines link data points collected on the same day from the same initial culture. Average cell volume of the culture was measured using a Coulter counter and normalized to the average cell size of a wild-type strain. (*WHI5*pr: n=9, *ACT1*pr: n=4) (C) Rpb1 ChIP-seq with spike-in normalization to measure RNA Pol II occupancy per cell for the indicated genes (data from Swaffer et al 2023^28^). Data were normalized to the average value for the indicated gene for wild-type and larger *cln3Δ* cells so they could be compared with our results in panel B. n=2 biological replicates. The line represents the linear regression taken from the four points for each gene. (D) Schematic representation of the various promoters whose size scaling properties were examined. The *WHI5* core promoter is defined as from -126 to -76bp upstream of the start codon, the *WHI5* 5’UTR is defined as from bp -75 to 0bp upstream of the start codon. The *ACT1* core promoter is defined as from -208 to -121bp, the *ACT1* 5’UTR is from -120 to 0 bp, and the *HIS3* core promoter is from -50 to -24bp. (E) Size scaling slope calculated as shown in panel C for the indicated promoters. Error bars indicate the 95% confidence intervals, and each data point represents a separate biological replicate. For the indicated comparisons, ns denotes p>0.05, * p<0.05, and ** p< 0.01. (545bp *WHI5*pr: n=9, *ACT1*pr: n=4, 136bp *WHI5*pr: n=8, 545bp *WHI5*pr[*ACT1*pr core+5’UTR]: n=5, 545bp *WHI5*pr[*ACT1*pr core]: n=5, 545bp *WHI5*pr[5’UTR]: n=4, *HIS3*pr: n=9, *HIS3*pr[*WHI5*pr core]: n=9) (F) mRNA concentrations expressed from the indicated promoter genotypes in small and large populations of cells (0 and 30ng/mL aTc, repectively). mRNA concentrations were measured relative to the endogenous *ACT1* mRNA. (545bp *WHI5*pr: n=9, *ACT1*pr: n=4, 136bp *WHI5*pr: n=8, 545bp *WHI5*pr[*ACT1*pr core+5’UTR]: n=5, 545bp *WHI5*pr[*ACT1*pr core]: n=5, 545bp *WHI5*pr[5’UTR]: n=4, *HIS3*pr: n=9, *HIS3*pr[*WHI5*pr core]: n=9) (G) The size scaling slopes of the Whi5-mCitrine protein synthesis rate calculated in single cells from time lapse imaging data as in Fig. 1D-E. For the indicated comparisons, ns denotes p>0.05, * p<0.05, and ** p< 0.01. (545bp *WHI5*pr: n=371, *ACT1*pr: n=169, 136bp *WHI5*pr: n=594, 545bp *WHI5*pr[*ACT1*pr core+5’UTR]: n=344, 545bp *WHI5*pr[*ACT1*pr core]: n=166)

Since measuring size scaling expression using RT-qPCR is relatively quick, we decided to proceed with this method to examine how a series of mutations to the *WHI5* promoter affected the size scaling of its expression (**Figure 2D**). We examined a truncated 136 base pair (bp) *WHI5* promoter and found that it also substantially sub-scaled with cell size similarly to the longer 545bp promoter (**Figures 2E and S2E**). Importantly, these changes to the scaling slope are not due to large changes in the expression level (**Figure 2F**). This led us to examine if either the 51bp core *WHI5* promoter or the 5’UTR was responsible for sub-scaling by swapping either one or both of these pieces with the equivalent sequences from the *ACT1* promoter. This experiment revealed that swapping out the 51 bp core *WHI5* promoter substantially reduced sub-scaling but that swapping the 5’UTR did not. We therefore conclude that the *WHI5* core promoter is required for the sub-scaling of *WHI5* expression. We next sought to test if the *WHI5* core promoter could promote sub-scaling gene expression in the context of another gene. To do this, we examined the effect of inserting *WHI5*’s core promoter in the place of the *HIS3* core promoter. While the expression of *HIS3* normally scales similarly to *ACT1*, this core promoter swap caused expression of *HIS3* to sub-scale (**Figure 2E**). However, a similar swap of the *WHI5* core promoter into the *ACT1* promoter did not lead to sub-scaling expression indicating that the ability of *WHI5*’s core promoter to promote sub-scaling expression depends on the context (**Figure S2F**). Taken together, this series of promoter mutations strongly argues that *WHI5*’s core promoter is largely responsible for its expression sub-scaling with cell size.

Having established that *WHI5*’s core promoter can drive sub-scaling mRNA concentrations using a series of qPCR experiments, we sought to verify this result using time lapse imaging experiments. Following our previously established methods^12^, we examined the rate of accumulation of Whi5-mCitrine fluorescent protein during S/G2/M phases of the cell cycle expressed from a series of promoters. When examining the size scaling of expression from an *ACT1* promoter, we only expressed the mCitrine fluorescent protein since the expression of a Whi5-mCitrine fusion protein from the *ACT1* promoter leads to very large cells^13^. These experiments measure the protein accumulation rate in cells of different sizes that are in precisely the same cell cycle phase. To expand the dynamic range of cell sizes, we again used an exogenously controlled *TETpr-WHI5* allele. These experiments corroborated our previous results and confirmed the critical importance of the core *WHI5* promoter for sub-scaling gene expression (**Figures 2G and S2G**).

### An array of Fkh transcription factor binding sites in the core *WHI5* promoter regulates sub-scaling gene expression

After identifying the 51bp *WHI5* core promoter as being largely responsible for its sub-scaling gene expression, we next sought to identify the underlying molecular mechanism. To do this, we first introduced 6 different linker scanning mutations to the *WHI5* core promoter where a 10 bp region was replaced with the 5’-TCTTACGATC-3’ sequence. In all cases, the size scaling of *WHI5* expression was increased, indicating that most parts of the core promoter contribute to sub-scaling (**Figures 3A and S3A**). A cursory examination of the core promoter sequence revealed an array of consensus or near consensus binding sites for the forkhead transcription factors Fkh1 and Fkh2^29,30^, which have been shown to bind Fkh1 and Fkh2 proteins in previous chromatin immunoprecipitation experiments^31^ (**Figures 3B and S3B**). To confirm this promoter binding, we tagged *FKH1* with a V5 tag and performed chromatin immunoprecipitation followed by qPCR to test for association with the *WHI5* promoter (see **methods**). This showed that Fkh1-V5 associated with the *WHI5* promoter and that this association was lost when the *WHI5* core promoter was swapped with the *ACT1* core promoter (**Figure 3C**). We performed an analogous experiment using *FKH2-V5* cells and found a similar *WHI5* core promoter association, but with a weaker signal (**Figure S3C**).

**Figure 3:**
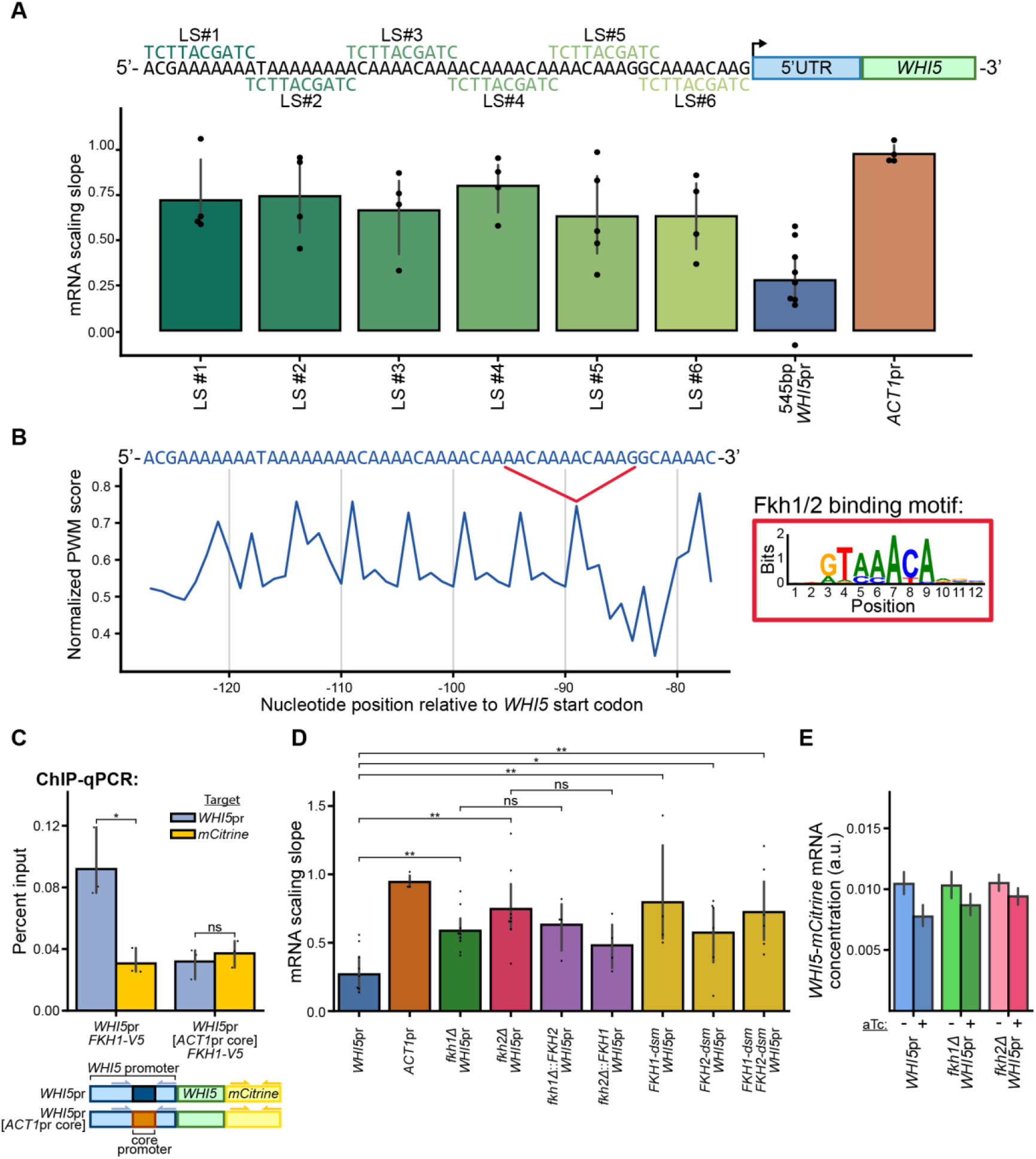
The forkhead transcription factors Fkh1 and Fkh2 bind an array of sites in the *WHI5* core promoter to regulate size sub-scaling expression. (A) Top: *WHI5* core promoter schematic indicating the 10bp regions that were substituted with the indicated sequences. Bottom: Size scaling slopes for mRNA expression calculated as in Fig. 2C shown for the indicated promoter mutants. We observed a p-value of <0.05 in student t-tests between the 545bp *WHI5*pr strain and each of the LS mutants. Error bars indicate 95% confidence intervals, and each data point represents a separate biological replicate. (545bp *WHI5*pr: n=9, all other strains: n=4) (B) Normalized position weight matrix score calculated using the Fkh-binding motif (logo shown top right). The 51 bp *WHI5* core promoter is shown in blue sequence. (C) Top: Chromatin immunoprecipitation followed by RT-qPCR performed for the indicated strains in which the endogenous *FKH1* gene has been fused to a V5 tag. Error bars indicate 95% confidence intervals, and each data point represents a separate biological replicate (n=3). Bottom: qPCR primers target the core promoter region or *mCitrine* (see **methods**). (D) Size scaling slopes for mRNA expression calculated as in **Figure 2C** shown for the indicated promoters with the indicated additional mutations. Error bars indicate 95% confidence intervals, and each data point represents a separate biological replicate. (*WHI5*pr: n=9, *ACT1*pr: n=4, *fkh1*Δ: n=10, *fkh2*Δ: n=9, *fkh1*Δ::*FKH2*: n=4, *fkh2*Δ::*FKH1*: n=4, *FKH1-dsm*: n=4, *FKH2-dsm*: n=6, *FKH1-dsm FKH2-dsm*: n=6) (E) mRNA concentrations expressed from the indicated promoter genotypes in small and large populations of cells (0 and 30ng/mL aTc, respectively). mRNA concentrations were measured relative to the endogenous *ACT1* mRNA. (*WHI5*pr: n=9, *fkh1*Δ: n=10, *fkh2*Δ: n=9)

That there were multiple Fkh binding sites in the *WHI5* core promoter and that Fkh proteins associated with the promoter suggested the hypothesis that Fkh1 and Fkh2 were responsible for sub-scaling *WHI5* expression. To test this hypothesis, we deleted *FKH1* or *FKH2* and then examined the size scaling of *WHI5* expression using RT-qPCR. Consistent with our hypothesis, we found a significant increase in the size-scaling of *WHI5* expression in both *fkh1Δ* and *fkh2Δ* cells (**Figure 3D**). This effect was not caused by changes in either the cell cycle distribution due to deletion of *FKH1* or *FKH2*, or large changes in *WHI5-mCitrine* expression levels (**Figures S3D and 3E**). Interestingly, the addition of an extra copy of *FKH1* to a *fkh2Δ* cell or the addition of an extra copy of *FKH2* to a *fkh1Δ* cell did not rescue sub-scaling. This suggests that both *FKH1* and *FKH2* are required for sub-scaling *WHI5* expression, possibly through interacting with one another. To test this, we decided to introduce mutations to the forkhead genes (*FKH1-dsm* or *FKH2-dsm*) that were shown to remove the ability of Fkh1 proteins to form a homodimer^32^. The *FKH2-dsm* mutation was not directly tested for its ability to disrupt dimerization, but the amino acid sequences of Fkh1 and Fkh2 are almost the same near the mutated residues^32^. This similarity between these two forkhead proteins in this region also suggests it might be used to form heterodimers. It was previously found that these mutations did not affect the ability of Fkh proteins to regulate transcription, but did affect the ability of Fkh proteins to mediate longer range genomic interactions likely through the two DNA-binding domains of the dimer^32,33^. That *FKH1-dsm* and *FKH2-dsm* cells both increased the size scaling of *WHI5* expression indicates that Fkh protein-protein interactions likely play a role in promoting sub-scaling gene expression.

The presence of an array of Fkh binding sites and the fact that Fkh proteins can interact with one another suggests a model where the array of binding sites promotes the cooperative binding of a Fkh protein filament to the core promoter region. In this model, a reduction in the number of binding sites should reduce the cooperativity of the binding and reduce the sub-scaling of *WHI5* expression. This is precisely what we observed in the 6 different linker scanning mutations to the *WHI5* core promoter (**Figure 3A**).

### A mathematical model shows cooperative Fkh binding can drive sub-scaling expression

Our results so far suggest a model where Fkh1 and Fkh2 transcription factors bind an array of Fkh sites on the *WHI5* core promoter to drive sub-scaling gene expression. One possible mechanism through which this might work would be through cooperative binding. In general, transcription increases in larger cells^7,34–38^. For most genes, the bulk of the increase in transcription is due to their increased nuclear concentration of free RNA polymerase II^28^. However, there are many other proteins and general transcription factors that accumulate on the genome in larger cells as can be seen in a chromatin enrichment proteomics experiment done on a population of larger and smaller cells^28^. Thus, there appears to be an accumulation of proteins on different promoter sequences that can promote the size scaling of gene expression. One way that *WHI5* could avoid the general size scaling of gene expression would be if the configuration of its promoter were more invariant with cell size. This could occur if the Fkh transcription factors bound cooperatively to the promoter so that it was fully occupied in both large and small cells.

To explore how Fkh binding to an array of sites on the *WHI5* core promoter could result in cooperative binding, we built a simple mathematical model (**Figure 4A**). For simplicity, we neglect the differences between Fkh1 and Fkh2 and model a single Fkh transcription factor that can form a filament as has been shown to be the case for the related forkhead transcription factor FOXP3 in mammalian cells^39^. In our model, the Fkh protein can both bind a target site on the genome and bind other Fkh proteins already bound to adjacent sites. We refer the reader to the **methods** for a complete model description. Since we do not know the free energies associated with any of these interactions, we simply model both as favorable binding events with a free energy change ΔG. Thus, a Fkh protein binding a target site on its own would correspond to a free energy change ΔG, while a Fkh protein binding a target site with two adjacent sites occupied by Fkh proteins would correspond to a three times as large free energy change 3ΔG. To complete the model, we assume the concentration of Fkh protein does not change with cell size, so that larger cells have proportionally more Fkh protein. Thus, in smaller cells, a higher proportion of Fkh protein is bound to the genome to reduce the concentration of the free nuclear Fkh protein, which in turn reduces the target site occupancy. In larger cells, which have more Fkh protein, the depletion of free nuclear Fkh protein from genome binding is much less pronounced so that its concentration is higher and the target site occupancy is increased (see **methods, Figure S4**). Importantly, the quantitative nature of these relationships depends on the particular binding parameters and concentrations in the model. However, the general trend that there should be a higher concentration of free transcription factor in larger cells capable of driving higher site occupancy is robust to parameter variation. Indeed, we recently reported a similar result when examining the occupancy of RNA Polymerase II on the budding yeast genome^28^. There, in a similar dynamic equilibrium model, we found that increased transcription in larger cells was driven by an increase in the free nuclear RNA Polymerase II concentration due to the proportionally larger number of these complexes in larger cells.

**Figure 4:**
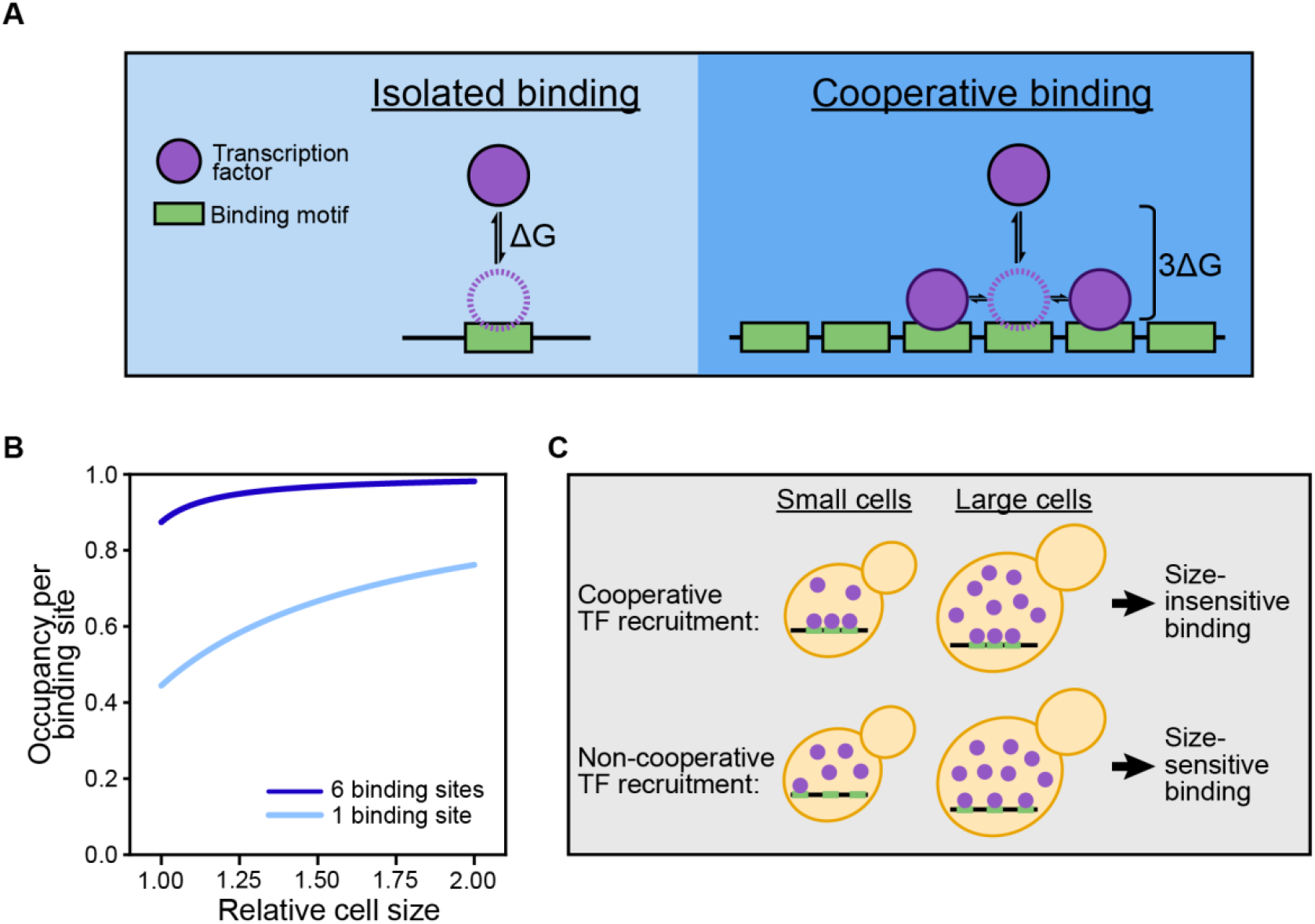
Cooperative binding can saturate an array of binding sites in smaller cells. (A) Schematic illustration of a mathematical model where a generic Fkh transcription factor can bind a site on the genome with a favorable free energy change ΔG. In a binding site array, Fkh can also bind adjacent bound Fkh proteins, also with a free energy change ΔG. In this case, the more Fkh proteins are bound, the more favorable it is to bind additional proteins, making the interaction between Fkh and the binding site array cooperative (see **methods**). (B) The Fkh concentration is constant so that larger cells have proportionally more Fkh proteins. In these larger cells, the depletion of free nuclear Fkh protein is much less pronounced so that its concentration is higher and the occupancy of single isolated Fkh sites increases. For an array of binding sites, which support cooperative binding, the sites are all occupied at smaller cell sizes. (C) Schematic representation of model results showing the increased occupancy of Fkh on single, isolated sites in larger cells, while an array of cooperatively-bound Fkh sites is occupied in both larger and smaller cells.

Having completed the specification of a simple binding model, we can now examine how the occupancy of a single Fkh binding site compares to an array of Fkh binding sites, where the free energy change is enhanced by the cooperative binding of adjacent Fkh proteins. For a typical set of parameter values, we see much higher occupancy of the site array compared to the single site. This results in the saturation of binding to the site array, while the binding of a single site continues to increase with cell size (**Figure 4B**). If Fkh-driven transcription is driven by site occupancy, then this could explain the relatively size-independent transcription of *WHI5*, while the transcription of other Fkh-target genes, which are mostly regulated by single or isolated sites, increases in proportion to cell size. Finally, we note the conclusions of this modeling work are tentative because we do not know the parameters. However, the general nature of cooperative binding models supports the qualitative conclusion that a multisite binding array will result in earlier saturation of the sites than would be the case for the less energetically favorable binding to a single site (**Figure 4C**).

### Structural models are consistent with Fkh1 and Fkh2 multimeric *WHI5* promoter binding

Our analysis so far suggests that a filament of Fkh1 and Fkh2 proteins binds an array of Fkh sites on the *WHI5* promoter to regulate its sub-scaling expression. To test the feasibility of multiple Fkh1 and Fkh2 proteins simultaneously occupying the *WHI5* core promoter, we sought to create predictive structural models using AlphaFold 3 ^40^. We tested two sets of input conditions: 3 Fkh1 proteins, 3 Fkh2 proteins, and the *WHI5* core promoter; and 2 Fkh1 proteins, 2 Fkh2 proteins, and the *WHI5* core promoter. We found that the output models are consistently generated with high pLDDT scores for the putative DNA binding domains (DBDs) of both Fkh proteins, suggesting high confidence in the predicted structures of the DBDs and their interaction with the core promoter DNA (**Figures 5A and S5A**). In contrast, other regions of both proteins had low pLDDT scores and were highly variable between models. We therefore focused our analysis on the DBDs and how they interact with DNA. Notably, DBDs of Fkh1 and Fkh2 bind to the tandem AAACA motifs in the core promoter without steric hindrance (**Figures 5B and S5B)**. In several models, Fkh1 and Fkh2 occupy consecutive consensus sites along the DNA such that all the Fkh1 promoters occupy one face of the DNA and the Fkh2 promoters occupy the opposite face (**Figure 5B**). In other models, Fkh1 or Fkh2 from two different heterodimers bind consecutive consensus sequences such that both proteins alternate along both faces of the DNA (**Figure S5B**). Importantly, all the models show an alpha helix from the DBD nestled in the major groove at each Fkh binding motif (**Figures 5B and S5B**). The model predicts that the asparagine and histidine residues (N349 and H353 in Fkh1, N386 and H390 in Fkh2) in this helix form hydrogen bonds with the nucleotides in the core promoter (**Figure 5C**). In particular, the histidine contacts the first adenine and complimentary thymine in the AAACA consensus, and the asparagine contacts the second adenine. These interactions are similar to those made by the DNA binding domains of the homologous mammalian transcription factors FoxP2 (PDB ID: 2a07)^41^ and FoxP3 (PDB ID:7tdw)^42^. Disrupting these interactions by mutating these residues to alanines in *FKH1* reduced the sub-scaling of *WHI5* expression (*FKH1-DBD*) (**Figure 5D**). There might also be an effect when a similar mutation is introduced into *FKH2*, but the effect size is likely much smaller (*FKH2-DBD*). These observations are consistent with a reduction in binding of the Fkh filament if one of the two Fkh proteins is no longer able to bind to the promoter. Taken together, our structural and mutational analysis indicate that a Fkh1-Fkh2 multimer could bind the *WHI5* core promoter.

**Figure 5:**
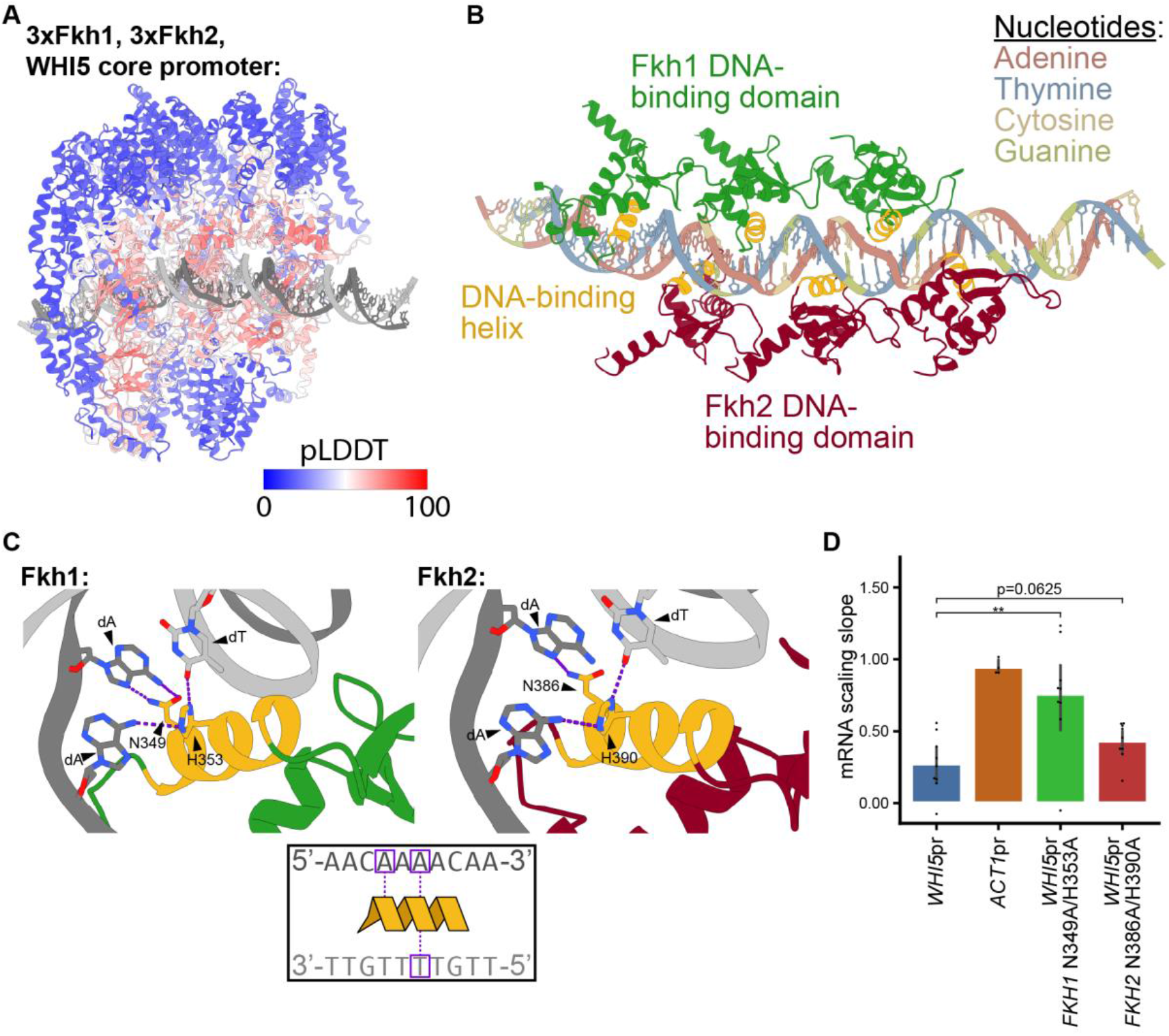
AlphaFold3 structural predictions are consistent with a Fkh1/2 filament binding an array of sites on the *WHI5* core promoter. (A) AlphaFold3 structural model of 3 Fkh1 proteins, 3 Fkh2 proteins, and the 51 bp *WHI5* core promoter. Output models consistently have high pLDDT scores for the putative DNA binding domains (DBDs) of both Fkh proteins, suggesting high confidence in the predicted structures of the DBDs and their interaction with the core promoter DNA. (B) DBDs of Fkh1 and Fkh2 bind to the tandem forkhead motifs in the core promoter, whose bases are color coded. An alpha helix from the DBD is nestled in the major groove at each forkhead binding motif. (C) The alpha helix’s asparagine and histidine residues (N349 and H353 in Fkh1, N386 and H390 in Fkh2) form hydrogen bonds with the indicated nucleotides (purple boxes in the inset) in the core promoter. (D) Size scaling slopes for mRNA expression calculated as in **Figure 2C** shown for the *WHI5* promoter with the indicated additional mutations. Error bars indicate 95% confidence intervals, and each data point represents a separate biological replicate. The Fkh1-DNA interaction is likely disrupted by mutating N349 and H353 to alanine (*FKH1-DBD*), and the Fkh2-DNA interaction is likely disrupted by mutating N386 and H390 to alanine (*FKH2-DBD*). (*WHI5*pr: n=9, *ACT1*pr: n=4, *FKH1* N349A/H353A: n=9, *FKH2* N386A/H390A: n=9)

### Proteomic analysis shows sub-scaling expression across the genome is not generally Fkh-dependent

Having established that Fkh1 and Fkh2 regulate the sub-scaling of *WHI5* expression, we sought to test if other sub-scaling genes were also regulated in this way. To do this, we examined how the proteome changes with increasing cell size in *fkh1Δ* and *fkh2Δ* cells compared with wild-type cells. We previously examined how protein concentrations changed in budding yeast cells using proteomics^23^. Here, we extend this analysis to populations of *fkh1Δ* and *fkh2Δ* mutant cells. To do this, we used our *TETpr-WHI5* strain to generate two populations of large and small cells for proteomics analysis (**Figures 6A and S6A**; see **methods**). For each protein that was reliably detected (**Figure S6B**), we calculated its relative concentration in small and large cells. From these relative concentrations, we calculated the size-dependence, or ‘size slope’ of each protein^23^. Replicate experiments gave highly reproducible results (**Figure S6C**). We then calculated how each gene’s size-scaling changed from that in wild-type cells in *fkh1Δ* or *fkh2Δ* strains. Interestingly, most proteins exhibited a similar size-dependence in wild-type, *fkh1Δ*, and *fkh2Δ* cells with only a small group of proteins changing upon *FKH* deletion (**Figure S6D**). In addition, we found that the originally sub-scaling genes were not more likely to become more scaling upon *FKH* deletion.

**Figure 6:**
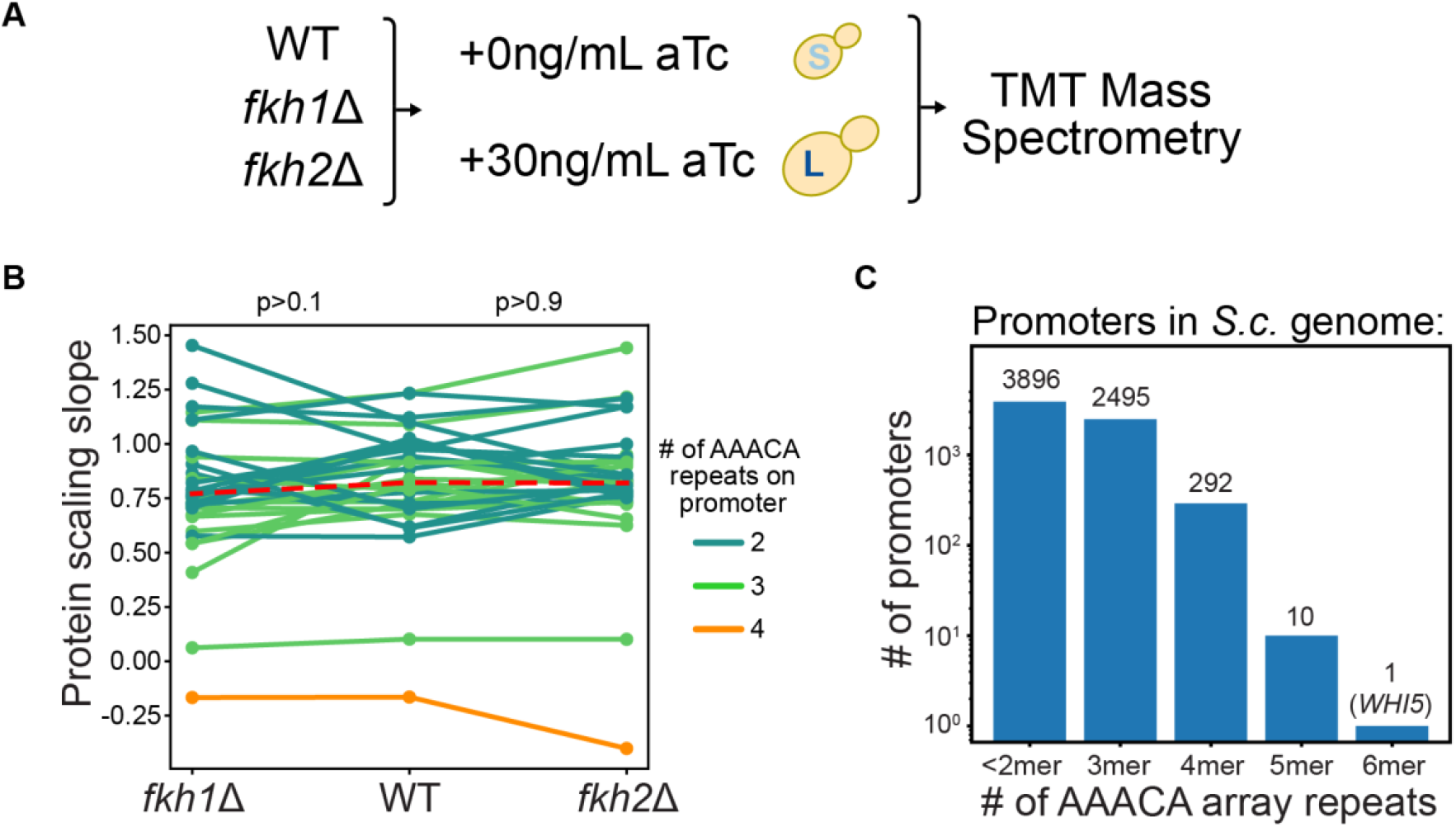
Proteomic and bioinformatic analysis highlights the uniqueness of the Fkh-binding site array driving sub-scaling *WHI5* expression. (A) Experiment schematic illustrating the workflow for mass spectrometry experiments examining how cell size impacts the proteome of wild-type, *fkh1Δ*, and *fkh2Δ* cells (see **methods**). As previously, an aTc-inducible *WHI5* allele is used to generate populations of small and large cells. (B) Scaling slopes for known Fkh target genes in wild-type, *fkh1Δ*, and *fkh2Δ* cells. Slopes for individual genes are connected by a line and the number of Fkh array binding sites in their promoters are denoted by color. (C) Number of Fkh concatemers within promoter sequences of all known ORFs in the budding yeast genome. Promoter sequences are scored with a concatenated Fkh 6-mer position weight matrix. The mean scores of random sequences with known n-mer motifs are then used to bin the distribution of scores.

Even though there was no global change in protein sub-scaling in *FKH* mutants, we sought to identify any shared promoter features in the small group of proteins that did increase their scaling slope. We took the subset of genes that increased their scaling slope upon *FKH* deletion by more than 2 standard deviations from the mean (83 genes for *fkh1Δ*, 69 genes for *fkh2Δ*) and performed *de novo* motif discovery on their promoter sequences (defined as 500bp upstream of the open reading frame) (**Figure S6D**). Surprisingly, we found no enrichment of Fkh binding motifs (**Figure S6E**). Furthermore, when we focused on the size-scaling of 35 previously identified Fkh1/2-regulated genes^43,44^, we saw no significant difference in their scaling upon *fkh1Δ* or *fkh2Δ* deletion (**Figure 6B**). Taken together, our proteomics analysis reveals that Fkh is likely not a major driver of global sub-scaling gene expression.

Since Fkh1 and Fkh2 had no global effect on protein sub-scaling, we examined features of *WHI5*’s promoter that may be unique. An analysis of budding yeast promoter sequences revealed *WHI5* has the highest number of repeated Fkh binding sites across the genome (**Figure 6C**). This suggests that the *WHI5* promoter may be uniquely specialized in using Fkh-heteropolymer-binding to regulate sub-scaling expression.

### Replacing *WHI5*’s core promoter reduces cell size control in G1

So far, we have shown that Fkh1 and Fkh2 regulate sub-scaling expression through *WHI5*’s core promoter. This sub-scaling expression contributes to the inverse correlation of Whi5 concentration and cell size in the G1 phase of the cell cycle^13^. To test if this mechanism contributes to cell size control at the G1/S transition, we sought to measure the degree of size control in cells where the *WHI5* core promoter was swapped with the *ACT1* core promoter and compare this with wild-type cells. We defined size control as the inverse correlation between how much daughter cells grew in G1 phase and how big they were when they were born^4,46^. To isolate the effect of the Whi5 size control pathway, we decided to perform this analysis in a strain background lacking the *BCK2* gene because it is responsible for an additional size control pathway^12,47–49^. Using time lapse imaging and analysis (see **methods**), we measured these quantities in single unperturbed cells that were growing and dividing asynchronously (**Figure 7A**). We found that the inverse correlation between the amount of growth in G1 and the cell size at birth was reduced in the core promoter swapped cells compared to wild-type cells, while the relationship between the size at budding (beginning of S phase) and the amount of growth in S/G2/M phases was unchanged (**Figures 7B-C**). These data showing reduced G1/S size control when the sub-scaling of Whi5 is reduced are consistent with a conceptually similar experiment where the entire *WHI5* promoter was replaced with the *TET* promoter and Whi5 was destabilized in order to reduce the importance of the chromatin-based partitioning mechanism^13^. Taken together, this series of experiments shows that G1 size control relies on the sub-scaling behavior of Whi5 protein concentration driven by the Fkh-regulated core *WHI5* promoter.

**Figure 7:**
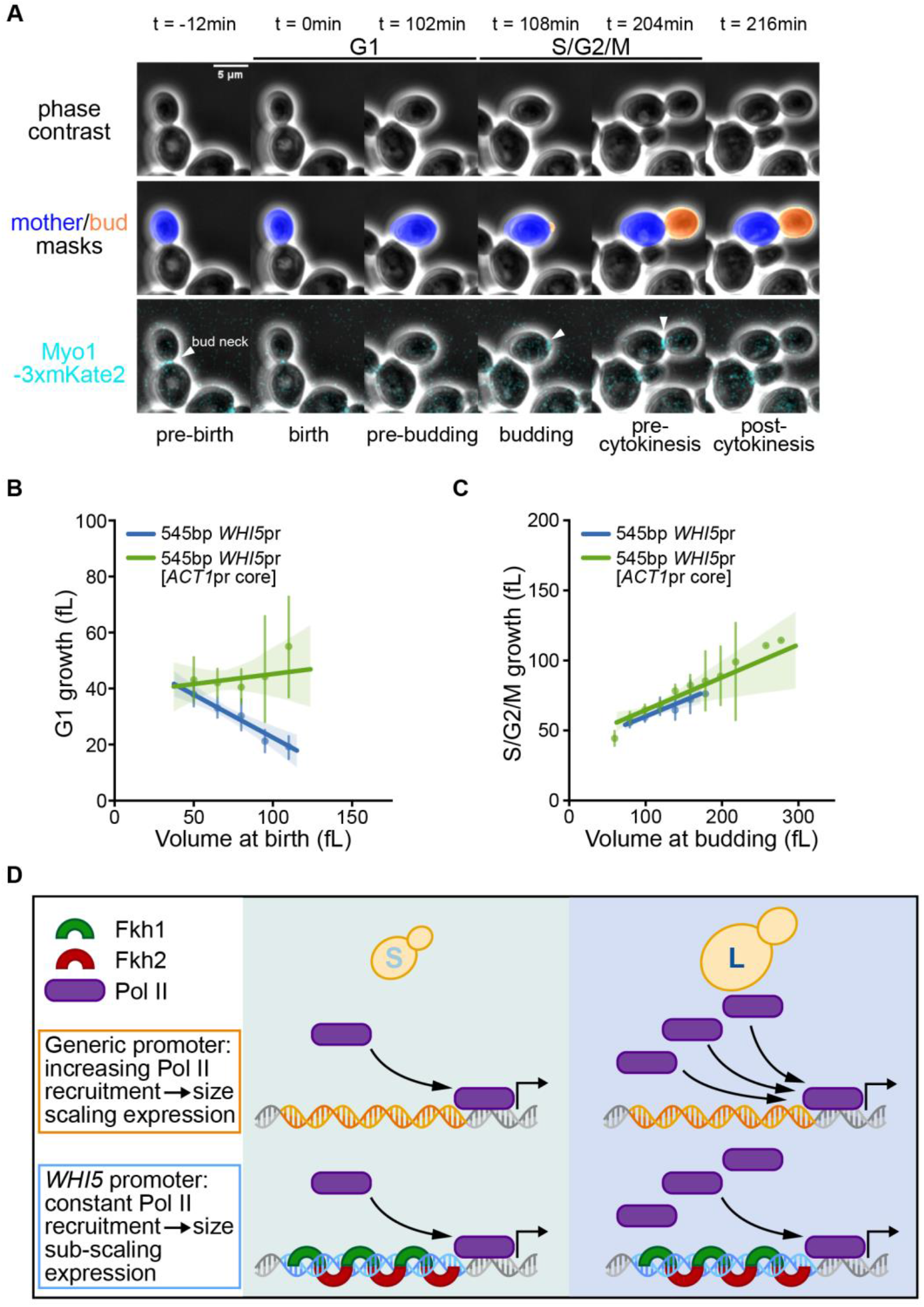
Replacing the *WHI5* core promoter weakens cell size control. (A) Top: phase contrast images of asynchronously dividing cells. Middle: Cells are segmented and tracked using the Cell-ACDC framework (see **methods**). Mother cells and their associated buds are tracked through entire cell division cycles. Bottom: Composite phase and fluorescence images showing the fluorescent Myo1 protein fused to 3 copies of the red fluorescent protein mKate2. Myo1 appears at budding and leaves the bud neck at cytokinesis to bookend the S/G2/M phases of the cell cycle. t=0 corresponds to the time of division of the indicated cell. (B) The amount of cell growth in the G1 phase of daughter cells plotted against the size of the daughter cell at birth for the indicated genotypes. Both strains contain *bck2Δ*. The lines denote linear fits. Error bars and shaded areas indicate 95% confidence intervals. (545bp *WHI5*pr: n=171, 545bp *WHI5*pr[*ACT1*pr core]: n=254) (C) The amount of growth in the S/G2/M phases of the cell cycle plotted against cell size at bud emergence, which marks the beginning of S phase. The lines denote linear fits. Error bars and shaded areas indicate 95% confidence intervals. (545bp *WHI5*pr: n=171, 545bp *WHI5*pr[*ACT1*pr core]: n=254) (D) Schematic diagram illustrating how an array of Fkh binding sites could regulate sub-scaling *WHI5* expression.

## DISCUSSION

A critical component of the budding yeast Whi5-dilution model is that the Whi5 concentration is inversely related to cell size at the beginning of G1 phase. This requires two mechanisms: the chromatin-mediated partitioning of Whi5 in nearly equal amounts at cell division and the sub-scaling of *WHI5* transcription in the S/G2/M phases of the cell cycle. Our study identifies the mechanism by which sub-scaling *WHI5* transcription remains largely independent of cell size, a property essential for robust size control at the G1/S transition. We show that this size-independent, sub-scaling expression is driven by the *WHI5* core promoter, which contains an array of Fkh1 and Fkh2 transcription factor binding sites. Genetic and structural perturbations to these forkhead factors and their binding motifs reduce the sub-scaling behavior of *WHI5* expression, supporting a model in which cooperative binding of a Fkh1/2 heteropolymer enables stable promoter occupancy regardless of cell size. This unique regulatory architecture allows *WHI5* transcription to resist the general scaling of transcription with cell volume, preserving the inverse relationship between Whi5 concentration and cell size at birth. By replacing the *WHI5* core promoter, we demonstrate that this transcriptional mechanism is important for effective size control during G1.

Since the Whi5 dilution model was first proposed over a decade ago, its validity has been the subject of some controversy^50–52^, which we have previously addressed in detail^4,14^. While part of the confusion stems from technical issues and interpretation, another important factor is that multiple size control mechanisms operate in all organisms that have so far been studied in detail^4,11,53–55^. As a result, disrupting a single mechanism does not necessarily lead to dramatic changes in cell size variability, but rather to more subtle phenotypes that require careful quantitative analysis. Our findings in this study contribute significantly to this discussion. We identify the molecular basis for the sub-scaling of *WHI5* transcription with cell size and show that disabling this mechanism reduces the size-dependence of the G1/S transition. These results solidify the central tenets of the Whi5 dilution model, namely, that Whi5 concentration decreases with increasing cell size in G1, and that elevated Whi5 concentrations in smaller cells delay the G1/S transition, thereby contributing to cell size control.

A key strength of this work is the identification of a molecular link between the sub-scaling expression of *WHI5* and the presence of a dense array of Fkh binding sites in its core promoter. Mutational analysis of this region showed that 10bp substitutions that remove subsets of Fkh sites lead to a reduction of sub-scaling expression, implicating cooperative Fkh binding in this regulation. Our structural modeling supports the feasibility of simultaneous binding by at least six Fkh1/2 proteins to the *WHI5* promoter. Cooperative binding would require interactions between these factors, which are known to occur in dimeric form^32^. Though direct evidence for higher-order heteropolymers is lacking, our structural modeling suggests there are multiple ways heterodimeric and homodimeric Fkh1/2 interactions could form while the protein DBDs are bound to the arrayed sites on the promoter. Interestingly, the mammalian forkhead protein FOXP3 forms higher-order multimers when engaging arrays of TnG repeats, which are very similar to those found in the *WHI5* core promoter^39^. Our genetic data show that deletion of either *FKH1* or *FKH2* cannot be compensated by overexpression of the other, suggesting a heteropolymeric assembly is required for this sub-scaling regulation, in contrast to the homopolymer observed for FOXP3.

This cooperative assembly of Fkh1/2 likely leads to saturation of RNA Polymerase II the *WHI5* promoter in both small and large cells, thereby decoupling transcription from cell size (**Figure 7D**). In contrast, other Fkh-regulated genes with less densely packed, or less cooperative Fkh binding sites, exhibit increased expression in larger cells, making *WHI5* unique among this cohort. Saturated Fkh binding at the *WHI5* promoter would render its occupancy insensitive to the global increase in general transcription factors on the genome as cells grow^28^, and thereby maintain a constant transcription rate in large and small cells.

More broadly, our results suggest that promoter architecture and transcription factor cooperativity can be finely tuned to control the scaling behavior of gene expression with respect to cell size. Importantly, whether a gene exhibits sub-scaling, scaling, or super-scaling behavior may depend not only on the identity of the transcription factors involved but also on how their binding sites are arranged within the promoter to generate cooperative interactions. As scaling behaviors of gene expression with cell size are widespread across eukaryotes, we anticipate that the Fkh-based mechanism we describe here may serve as a paradigm for other systems in which precise size-dependent control is critical to cellular function.

## Acknowledgements

This work was supported by the NIH through the R35 GM134858 award to JMS, the R35 GM145255 award to SMR, and the T32 training grant T32GM136631. We thank Blair Schooling and Masaru Shimasawa for assistance with microscopy experiments and Gheorghe Chistol for helpful discussions.

## Author contributions

Jacob Kim contributed to all aspects of this work, Shicong Xie performed bioinformatic analysis, Mike Lanz performed mass spectrometry experiments and analysis, Xin Gao, Jordan Xiao, and Kurt Schmoller performed and analyzed imaging experiments, Lucas Fuentes Valenzuela performed mathematical modeling, Corbin Mitchell and Seth Rubin performed structural analysis, Matthew Swaffer contributed to experimental design and data from his previous publication, and Jan M. Skotheim supervised the work and co-wrote the manuscript.

## STAR METHODS

### Key resources table

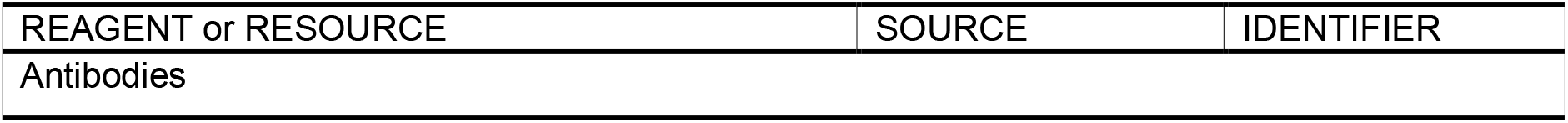
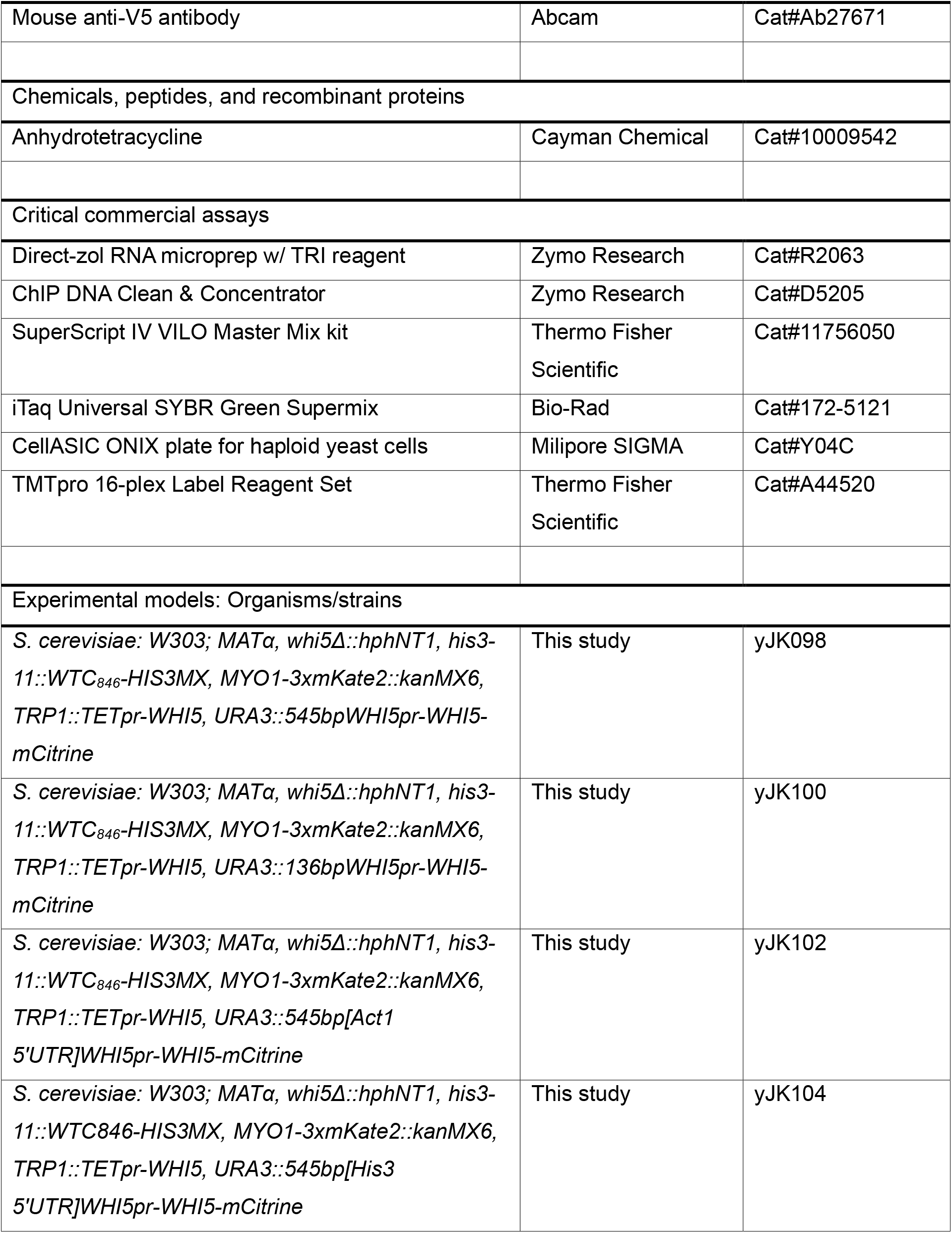
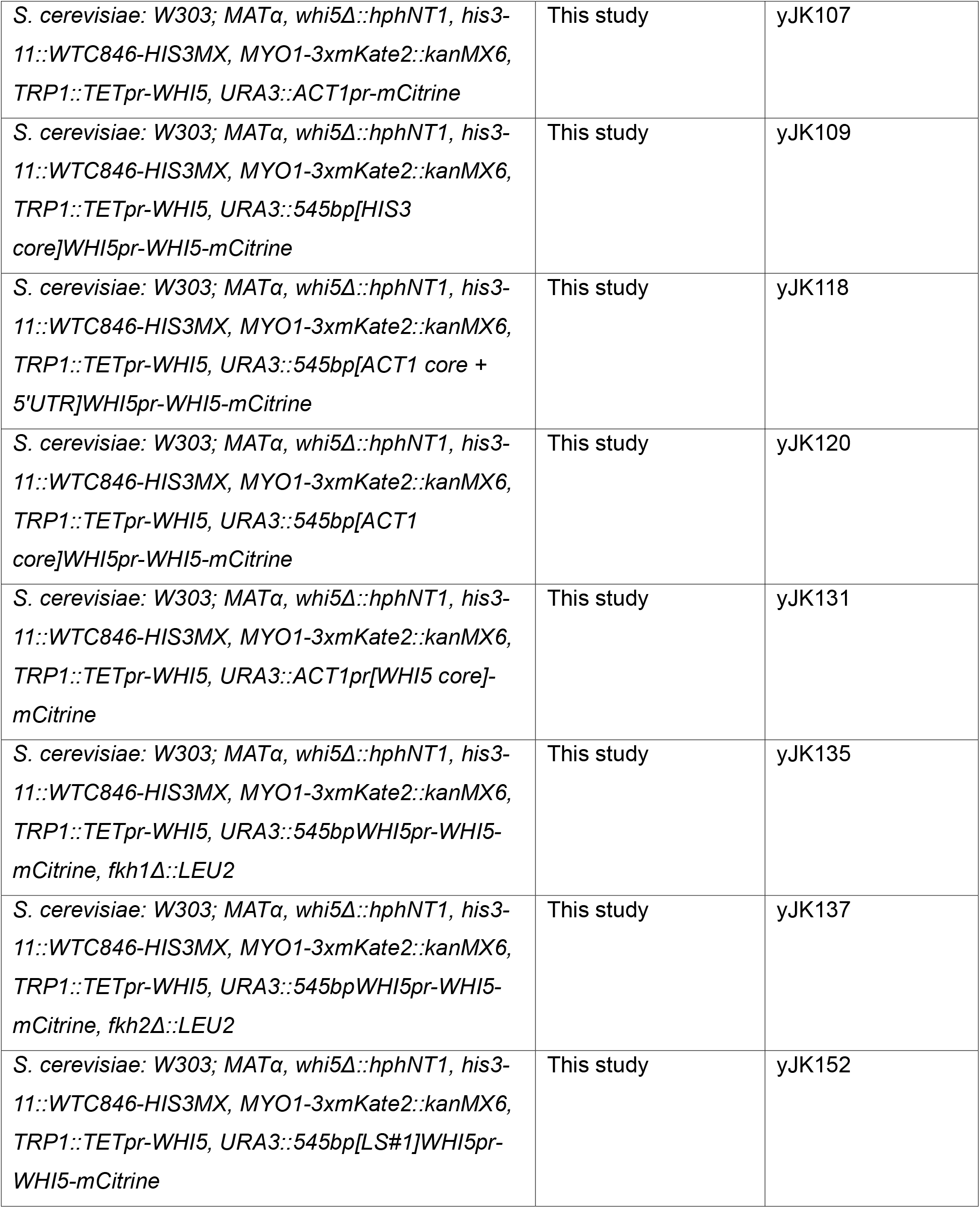
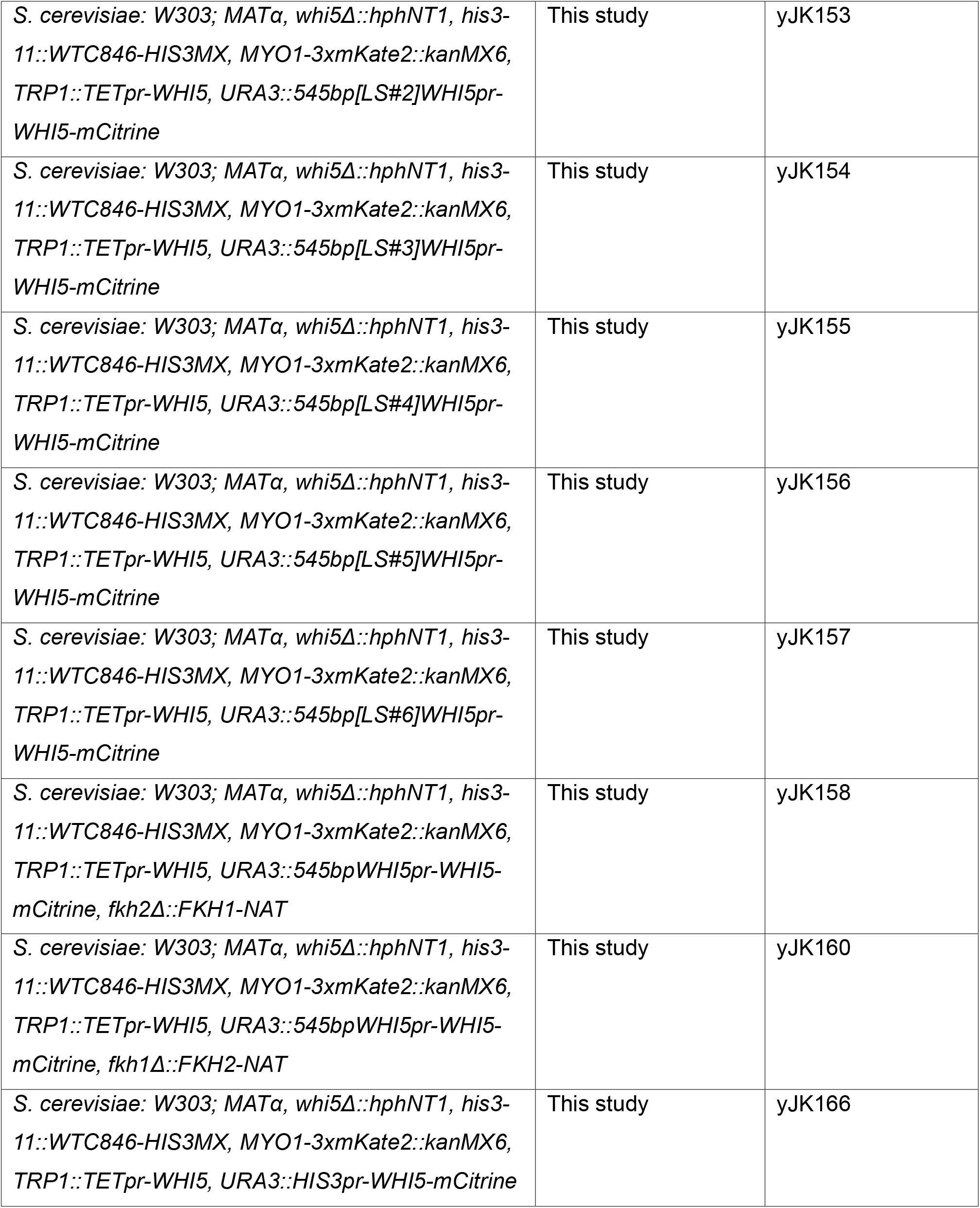
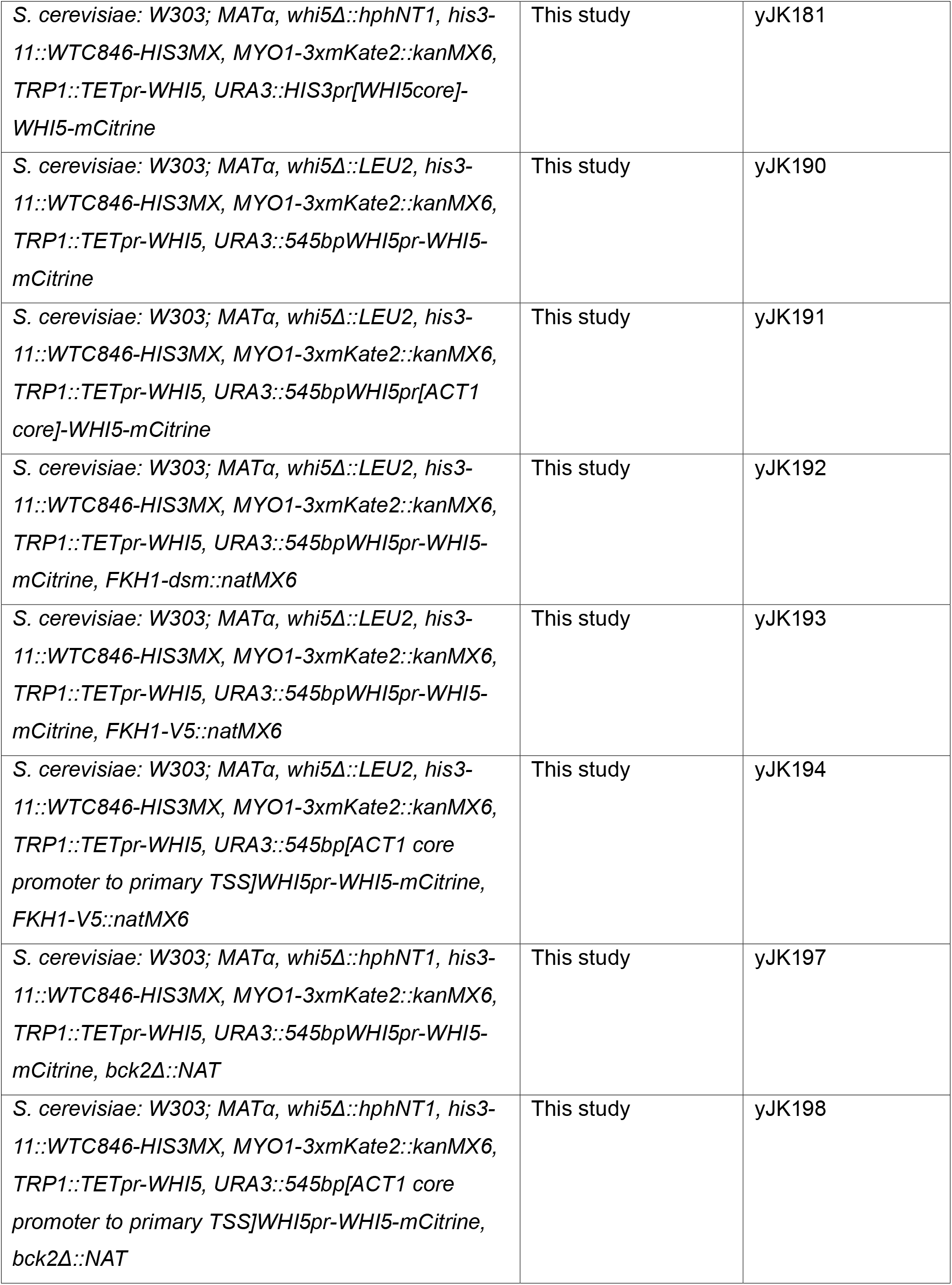
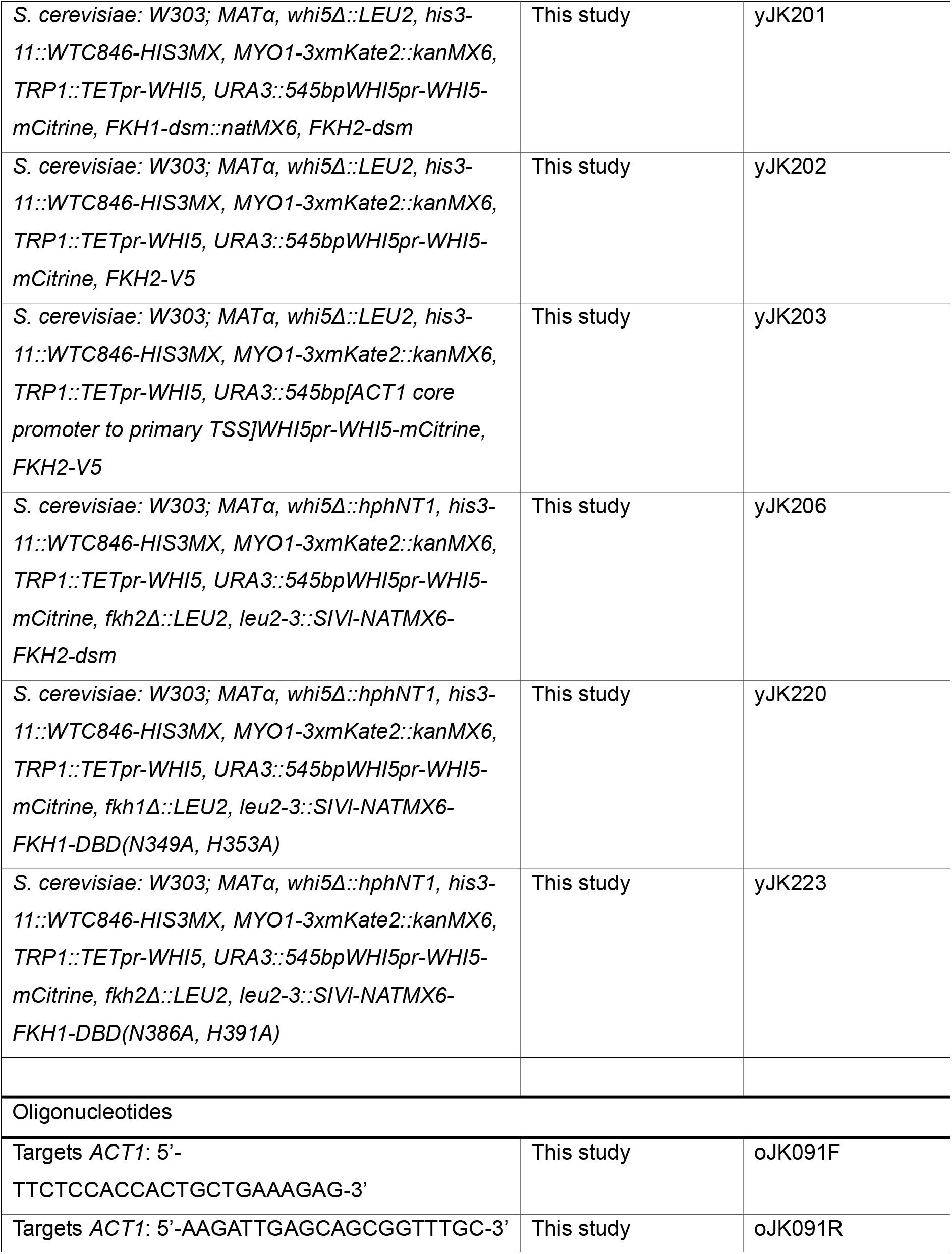
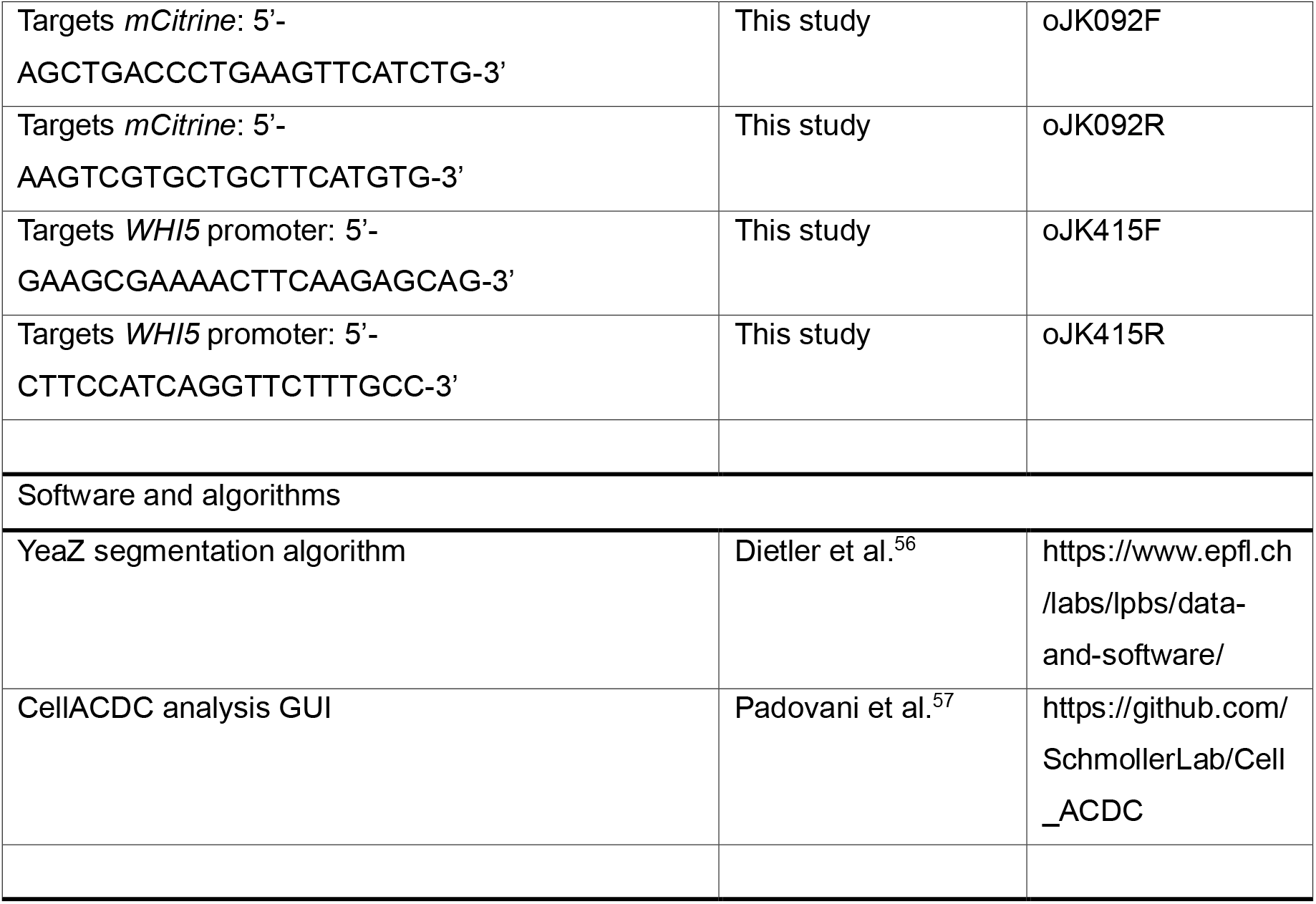

### Resource Availability

#### Lead contact

Further information and requests for resources and reagents should be directed to and will be fulfilled by the lead contact, Jan Skotheim.

#### Materials availability

All plasmids and strains generated in this study are available from the lead contact upon request without restriction.

#### Data and Code Availability

Bioinformatic and proteomic analysis code are available on the Skotheimlab GitHub. Raw images and image data are available upon request.

### Experimental model and study participant details

Standard procedures were used for growth and genetic manipulation of *Saccharomyces cerevisiae*. Full genotypes of all strains used in this study are listed in the methods and the Key Resource Table. All strains used were derived from the W303 background. However, the *ACT1* core promoter and 5’ UTR were determined from the genome of reference strain S288C. Unless otherwise indicated, cells were grown at 30°C in synthetic complete medium with 2% glycerol and 1% ethanol (SCGE).

### Method details

#### Microscopy

Microscopy experiments were performed similarly to previously^58^. Cells were first grown to be in log-phase (OD600 0.08-0.2) in SCGE media, then lightly sonicated before being loaded into a CellASIC Y04C microfluidics plate (Millipore SIGMA) under continuous SCGE media flow at 2psi. Images were taken every 6 minutes using an Observer Z1 microscope equipped with an automated stage, a plan apochromat 63x/1.4NA oil immersion objective, and an Axiocam 705 mono camera. Phase contrast images were taken using a HAL 100 halogen lamp at 5V power and 50ms exposure. Whi5-mCitrine was imaged using 400ms exposures under illumination from a Colibri 505nm LED module at 25% power. Myo1-3xmKate2, used to determine timing of cytokinesis, was imaged by exposure for 250-1000ms using a Colibri 555nm LED module at 25% power. Cells grown in the microfluidics plate were checked for their average growth rate to ensure it was similar to that in liquid culture. For each imaging experiment, we performed at least two biological replicate experiments, *i*.*e*., we examined separately inoculated cultures on different days.

#### Microscopy analysis synthesis rate estimates

Phase contrast images were segmented and tracked using the YeaZ convolutional neural network and tracker^56^ and manual correction through the Cell-ACDC framework^57^. Cell volume was estimated by finding the segmented 2D area’s longest axis, drawing a circle at each 1D slice perpendicular to the longest axis, and summing the areas of the circles together. To quantify relative mCitrine amounts, we first corrected for non-cell background fluorescence using a set of dark frame and flat field images. Raw images were initially corrected by subtracting the baseline signal present in the absence of illumination (dark frame). Then pixel values were rescaled using a cell-free median image (flat field) to account for reproducible spatial variations across the imaging field. Cell autofluorescence was corrected by subtracting a volume-dependent baseline determined from a linear fit to data from cells lacking fluorescence in the relevant channel. Total fluorescence was calculated as the sum of pixel intensities within the segmented cell area, minus the non-cell background signal and volume-dependent autofluorescence.

Mother–bud pairs were assigned in Cell-ACDC, with Myo1-3xmKate2 signal appearance and disappearance used to determine the timing of budding and cytokinesis, respectively. Unless otherwise indicated, all analyses of cell size control included only first-generation daughter cells with complete cell cycle traces. Mother cells do not exhibit cell size control and were not used in the analysis^46^.

To estimate Whi5-mCitrine and mCitrine synthesis rates, we analyzed time series for the amount of Whi5-mCitrine in the S/G2/M phase of the cell cycle (the budded phase). We fit a linear regression line to the data in S/G2/M and took the slope as an estimate of the Whi5 synthesis rate because Whi5 is a highly stable protein^12^. Synthesis rate estimates for individual cells were plotted against the cell size at the time of bud emergence, with both axes normalized by their respective mean value. The size-scaling slope was calculated using the slope of the linear regression fitted to this plot.

#### Flow cytometry measurements of DNA content

Cells were grown to log phase in SCGE for 48 hours. At an OD_600_ of 0.3, 1mL of absolute ethanol was added to 0.5mL of cells and the samples were placed in 4°C. The samples were then centrifuged at 4°C at 5500g for 5 minutes and the supernatant was removed. Then, 0.5mL syringe-filtered 50nM sodium citrate (pH 7.5) was added to pellet and incubated at room temperature for 30 minutes. The cells were centrifuged at 4°C at 10000g for 1 minute, and the supernatant was replaced with 0.5mL sodium citrate solution with 0.25mg/mL RNase A. After incubating for 1 hour at 55°C, 0.1mg proteinase K was added and incubated at 55°C for another hour. 8µg propidium iodide (PI) was added, and the samples were stored at 4°C in the dark overnight. The samples were vortexed and sonicated and placed on ice in the dark before flow cytometry analysis. Fluorescence from PI was measured for 10,000 events on an Attune NxT flow cytometer. After gating for single cells using the FlowJo software, a histogram of PI fluorescence intensity was plotted and a threshold was set just after the 1C peak. The G1 fraction was determined as the fraction of cells below the PI intensity threshold.

#### RT-qPCR

Cells were grown in SCGE with or without 30ng/mL aTc to 0.3 OD_600_. Cells were collected separately for size measurement using the Beckman Coulter Z2 Cell and Particle Counter and pelleting. 1-1.5mL of the cultures were centrifuged at 4°C at 21130g for 30 seconds, snap frozen in liquid nitrogen, and stored in -80°C for no more than 1 month. Frozen cell pellets were resuspended in TRI Reagent (Zymo Research) then lysed by bead beating using an MP Bio FastPrep-24 (MP Biomedicals, Santa Ana, CA; settings: 5.0m/s, 1 x 35 seconds) in a cold room at 4°C. Cell debris was pelleted by centrifugation (17,000g, 5 minutes) and the supernatant recovered. RNA was then extracted using a Direct-Zol RNA microprep kit (Zymo Research). Complementary DNA (cDNA) was synthesized using the SuperScript IV VILO synthesis kit (Thermo Fisher Scientific) on 300ng RNA. The cDNA was diluted 1:5 in Tris EDTA buffer, and stored in -20°C. The cDNA was further diluted 1:10 in water before using 4µL in each qPCR reaction. qPCR reactions were performed using iTaq Universal SYBR Green Supermix (Bio-Rad).

#### Measurement of mCitrine mRNA per cell

RT-qPCR measurements of target mRNA/*ACT1* mRNA (2^ΔCt^) were multiplied by the mean volume measured from the Beckman Coulter Z2 Cell and Particle Counter. Since *ACT1* mRNA scales very well with cell size and therefore is used frequently as a housekeeping gene, multiplying the 2^ΔCt^ value by the mean volume would result in relative target mRNA amounts per cell.

#### Chromatin Immunoprecipitation qPCR (ChIP-qPCR)

ChIP was performed similarly to previously^28^. 100mL cultures were grown in SCGE to 0.4 OD_600_, fixed for 20 minutes by adding formaldehyde to a final concentration of 1%. Cells were then quenched with glycine (final concentration 0.125M) for 5 minutes, washed twice in cold PBS, split equally into two tubes, pelleted, snap-frozen, and stored at -80°C. Pellets were thawed and lysed in 300μl FA lysis buffer (50mM HEPES–KOH (pH 8.0), 150mM NaCl, 1 mM EDTA, 1% Triton X-100, 0.1% sodium deoxycholate, 1mM PMSF, Roche protease inhibitor) with ∼1ml ceramic beads on a Fastprep-24 (MP Biomedicals). The entire lysate was then collected and adjusted to 1 ml before sonication with a 1/8” microtip on a Q500 sonicator (Qsonica) for 12 min (cycles of 10 seconds on and 20 seconds off). The sample tube was held suspended in a −20°C 80% ethanol bath to prevent sample heating during sonication. Cell debris was then pelleted, and the supernatant was retained for ChIP (985μL) or input (15μL). For each ChIP reaction, 20μl Protein G Dynabeads (Invitrogen) were blocked (PBS + 0.5% BSA, incubate 30-45 min at room temperature), pre-bound with 1μl of Mouse anti-V5 antibody (Abcam ab27671) in PBS (incubate 30-60 min at room temperature), and washed 2x with PBS before being incubated with supernatant (4°C overnight). After overnight incubation, Dynabeads were washed (5min per wash) 2x in FA lysis buffer and 3x in high-salt FA lysis buffer (50 mM Hepes-KOH (pH 8.0), 500 mM NaCl, 1mM EDTA, 1% Triton X-100, 0.1% sodium deoxycholate, 1mM PMSF). ChIP DNA was then eluted in ChIP elution buffer (50mM Tris-HCl (pH 7.5), 10mM EDTA, 1% SDS) at 65°C for 20 min. Eluted ChIP DNA or input DNA were then incubated to reverse crosslinks (65°C, O/N) before treatment with RNAse A (37°C, 1h) and then Proteinase K (65°C, 2h). DNA was then purified using the ChIP DNA Clean & Concentrator kit (Zymo Research). To ensure the size distribution of the sheared chromatin was similar, the input samples were analyzed using Bioanalyzer High Sensitivity DNA (Agilent). The DNA was diluted 1:10 before using 4μL per qPCR reaction.

#### Fkh cooperativity modeling

We constructed a model for the cooperative binding of Forkhead (Fkh) transcription factors to an array of sites arranged linearly (**Figure 4A**). For simplicity, we assume that Fkh1 and Fkh2 are the same and consider only one Fkh transcription factor that can form filaments. Each binding site can either be unoccupied or bound to a molecule from solution. Binding and unbinding are reversible, governed by the kinetic on and off rates, *k*_on_ [*C*] and *k*_off_, where [*C*] is the concentration of free Fkh molecules in solution. We first consider isolated binding of Fkh to a single site on the genome. At equilibrium, the ratio of these rates defines the dissociation constant 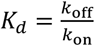. This equilibrium constant is related to the standard Gibbs free energy of binding Δ*G* = *RT* In 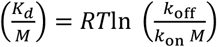, where M is the molar concentration unit, and *R* is the gas constant. We can solve this for the unbinding rate so that *k*_off_ = *k*_on_ M ⋅ *e*^*β*Δ*G*^, where 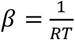 In our model, cooperative effects arise from neighboring bound molecules. Here, we take Δ*G* to be the free energy change associated with Fkh binding an isolated target site and ΔΔ*G* to be the additional free energy change associated with interacting with an already bound Fkh factor at a neighboring site. For simplicity, we will assume ΔΔ*G* = Δ*G*. Here, we take *n* to be the number of occupied neighbor sites so that the total free energy of binding at a site is Δ*G*(*n*) = Δ*G* + *n* ⋅ Δ*G* so that the effective off rate when there are n neighbors bound is:

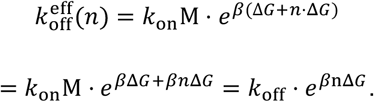

Since Δ*G* < 0, the presence of neighbors stabilizes binding and decreases the unbinding rate exponentially.

To estimate the average site occupancy, we use a mean-field approximation. We assume each neighbor is independently occupied with probability *p*, so the number of occupied neighbors *n* follows the binomial distribution: *P*(*n* = 0) = (1 − *p*)^2^; *P*(*n* = 1) = 2*p*(1 − *p*); *P*(*n* = 2) = *p*^2^. If we take *x* = *e*^*β*Δ*G*^, then the average unbinding factor becomes:

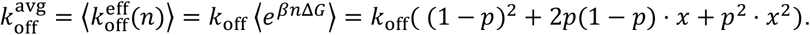

At equilibrium, the rate of binding to a site equals the rate of unbinding so that:

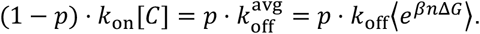

To simplify the equation, we use the dimensionless concentration 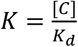 and get:

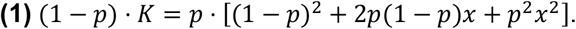

This nonlinear equation can be solved numerically to estimate the steady-state occupancy.

For a finite chain with *N* sites and open boundaries, the two edge sites have only one neighbor. We average the neighbor-dependent term accordingly:

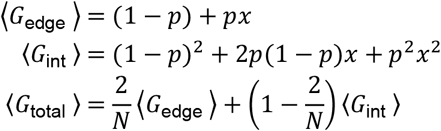

Then the corrected mean-field equation **(1)** becomes:

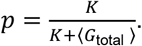

Results of the analytical model are given in **Figure 4B**. We see two limiting behaviors appearing. First, when the Fkh concentration *K* is small, the unbinding rate is much larger than the binding rate. In this case, the occupancy is always low and cooperative binding has little impact. Second, when *K* is large, the unbinding rate is much smaller than the binding rate so that sites are generally occupied and this behavior is enhanced by cooperative binding. Finally, when *K* ∼ 1, we see a strong transition from a small number of sites to a larger number of sites. The cooperative binding between neighboring sites strongly influences the mean occupancy (**Figure 4**).

The free Fkh concentration can increase with cell size if background binding sites are being diluted so that a smaller fraction of the total number of Fkh molecules are bound to the genome. We are assuming that the total Fkh concentration is constant as cells (and their nuclei) grow larger. In this case, the proportion of the Fkh molecules are bound to the genome decreases as cells grow larger, which results in a higher concentration of free Fkh molecules. Mathematically, this is equivalent to *k*_*on*,1_[*C*_*fkh,free*_][*C*_*sites, free*_] = *k*_*off*,1_[*CX*], where *k*_*on*,1_ and *k*_*off*,1_ are the binding rate of Fkh to background sites, [*C*_*fkh,free*_] and [*C*_*sites, free*_] are the free Fkh and background site concentrations, respectively, and [*CX*] is the bound site concentration. This can be rewritten as:

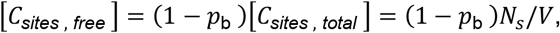

where *N*_*s*_ and *V* denote the number of sites and the cell volume, respectively, and *p*_b_ is the probability that a binding site is occupied by a Fkh molecule. On the other hand, we have that

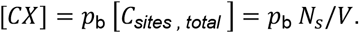

We can now use the fact that the total concentration of Fkh *C*_0_, is constant so that [*C*_*fkh,free*_] + [*CX*] = *C*_0_ and [*C*_*fkh,free*_] = *C*_0_ − *p*_b_ *N*_*s*_/*V*. Rearranging yields *k*_*on*,1_(*C*_0_ − *p*_*b*_*N*_*s*_/*V*)(1 − *p*_*b*_)*N*_*s*_/*V* = *k*_*off*,1_*p*_*b*_*N*_*s*_/*V*, which is equivalent to (*C*_0_ − *p*_*b*_*N*_*s*_/*V*)(1 − *p*_*b*_) = *K*_*D*,1_*p*_*b*_. This can be rearranged to find

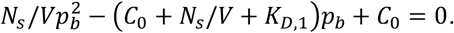

Solving this equation yields the implicit dependence of the binding probability on the cell volume, yielding the free Fkh concentration as a function of cell volume. As shown in **Figure S4**, under realistic parameters, the free concentration of Fkh increases nearly-linearly with cell volume. This justifies the approximation taken in **Figure 4B**.

Taken together, our mean-field formulation provides a simple, analytical way to estimate the average site occupancy for an array of Fkh binding sites with local cooperative effects. We see that a transition appears when the free Fkh concentration is similar to the dissociation constant (*K* ∼ 1), where we see a strong impact of cooperative binding on the mean occupancy. Those results are valid under the assumption that the change in free energy associated to cooperative binding is similar in magnitude to the natural change in free energy associated to binding.

#### Proteomics Sample Preparation

Cleared yeast lysates were alkylated with 5mM iodoacetamide and then precipitated with three volumes of a solution containing 50% acetone and 50% ethanol. Proteins were re-solubilized in 2M urea, 50mM Tris-HCl, pH 8.0, and 150mM NaCl, and then digested with TPCK-treated trypsin (50:1) overnight at 37°C. Trifluoroacetic acid and formic acid were added to the digested peptides for a final concentration of 0.2%. Peptides were desalted with a Sep-Pak 50mg C18 column (Waters). The C18 column was conditioned with 500µl of 80% acetonitrile and 0.1% acetic acid and then washed with 1000µl of 0.1% trifluoroacetic acid. After samples were loaded, the column was washed with 2000µl of 0.1% acetic acid followed by elution with 400µl of 80% acetonitrile and 0.1% acetic acid. The elution was dried in a Concentrator at 45°C. De-salted peptides were resuspended in 0.1% Formic acid. 25µgs of desalted peptide samples were resuspended in 20µLs of 100mM Triethylammonium bicarbonate solution and labeled with 16-plex TMTpro at a ratio 4:1 (TMT:peptide). Total reaction volume was less than 25µLs. The labeling reaction was quenched with a final concentration of 0.5% hydroxylamine for 15 min. Labeled peptides were pooled and acidified to pH ∼2 using drops of 10% TFA. Excess TMT label was removed by re-running the pooled sample through a Sep-Pak 50-mg C18 column (as described above).

#### Proteomics data acquisition and search

Data acquisition was performed as described previously^23^. TMT-labeled peptides were resuspended in 0.1% formic acid analyzed on a Fusion Lumos mass spectrometer (Thermo Fisher Scientific, San Jose, CA) equipped with a Thermo EASY-nLC 1200 LC system (Thermo Fisher Scientific, San Jose, CA). Peptides were separated by capillary reverse phase chromatography on a 25 cm column (75µm inner diameter, packed with 1.6µm C18 resin, AUR2-25075C18A, Ionopticks, Victoria Australia). Peptides were introduced into the Fusion Lumos mass spectrometer using a 180-min stepped linear gradient at a flow rate of 300nl min−1. The steps of the gradient are as follows: 6–33% buffer B (0.1% (v:v) formic acid in 80% acetonitrile) for 145 min, 33-45% buffer B for 15 min, 40–95% buffer B for 5 min and maintain at 90% buffer B for 5 min. The column temperature was maintained at 50 °C throughout the procedure. Xcalibur software (v.4.4.16.14) was used for the data acquisition and the instrument was operated in data-dependent mode. Advanced peak detection was disabled. Survey scans were acquired in the Orbitrap mass analyzer (centroid mode) over the range 380– 1,400 m/z with a mass resolution of 120,000 (at m/z 200). For MS1, the normalized AGC target (%) was set at 250 and maximum injection time was set to 100ms. Selected ions were fragmented by collision-induced dissociation (CID) with normalized collision energies of 34 and the tandem mass spectra were acquired in the ion trap mass analyzer with the scan rate set to ‘Rapid’. The isolation window was set to the 0.7-m/z window. For MS2, the normalized AGC target (%) was set to ‘Standard’ and maximum injection time to 35ms. Repeated sequencing of peptides was kept to a minimum by dynamic exclusion of the sequenced peptides for 30 s. The maximum duty cycle length was set to 3 s. Relative changes in peptide concentration were determined at the MS3 level by isolating and fragmenting the five most dominant MS2 ion peaks.

All raw files were searched using the Andromeda engine embedded in MaxQuant (v2). Reporter ion MS3 search was conducted using TMTpro (16-plex) isobaric labels. Variable modifications included oxidation (M) and protein N-terminal acetylation. Carbamidomthyl (C) was a fixed modification. The number of modifications per peptide was capped at five. Digestion was set to tryptic (proline-blocked). Database search was conducted using the UniProt Yeast proteome. The minimum peptide length was 7 amino acids. 1% FDR was determined using a reverse decoy proteome.

#### TMT proteomics analysis

Our TMT analysis pipeline utilized the peptide feature information in MaxQuant’s “evidence.txt” output file. Each row of the evidence.txt file represents an independent peptide and its corresponding MS3 reporter ion measurements. Peptides without signal in any of the TMT channels were excluded. Peptide measurements were assigned to a protein based on MaxQuant’s ‘Leading razor protein’ designation. For each individual peptide measurement (i.e., each row in the evidence table), the fraction of ion intensity in each TMT channel was calculated by dividing the ‘Reporter ion intensity’ column by the sum of all reporter ion intensities. To correct for loading differences between the TMT channels, each reporter ion channel was then normalized by dividing the fraction of ion intensity in each channel by the median fraction for all measured peptides (i.e., the median value for each column). This normalization scheme ensures that each individual peptide measurement is equally weighted when correcting for loading error.

#### Quantification of protein size-scaling

To quantify the proteome fraction (*i*.*e*. concentration) of each protein, the normalized ion intensity of all peptides identified across each protein was summed to obstrain a normalized protein intensity. For each protein, an average normalized protein intensity was obtrained as the mean across two independent replicates. To obtain the size-scaling slope, we calculated the ratio between the protein concentration in small and large cells (0 and 30ng/mL aTc, respectively) cells for WT, *fkh1Δ*, and *fkh2Δ*. Data is available in **Supplemental Table 1**.

#### AlphaFold analysis

AlphaFold3 models of Fkh1-Fkh2-DNA complexes were generated using the AlphaFold server through its web interface (https://alphafoldserver.com). Full-length Fkh1 and Fkh2 sequences were entered with either two or three copies each. A DNA sequence from the core Whi5 promoter was used (5’-ACGAAAAAAATAAAAAAAACAAAACAAAACAAAACAAAACAAAGGCAAAAC-3’).

#### Bioinformatic analysis of S. cerevisiae promoters

To find Fkh repeat arrays, we used the position weight matrix of Fkh1/2 to score the promoter sequence 500bp upstream of all putative ORFs in the *S. cerevisiae* reference genome. Only the high-consensus positions of the Fkh1/2 matrices corresponding to AAACA were used^59–61^. We concatenated multiple copies of the matrix and calculated each promoter’s score as a function of concatemer number. To discover *de novo* motifs that are enriched in genes whose expressions change upon *fkh1Δ* or *fkh2Δ* deletion, we used the XSTREME suite, which combines multiple motif discovery tools, using default settings: *E < 0*.*05*; 6 ≤ motif width ≤ 15^62^.

## SUPPLEMENTAL FIGURES

**Figure S1:**
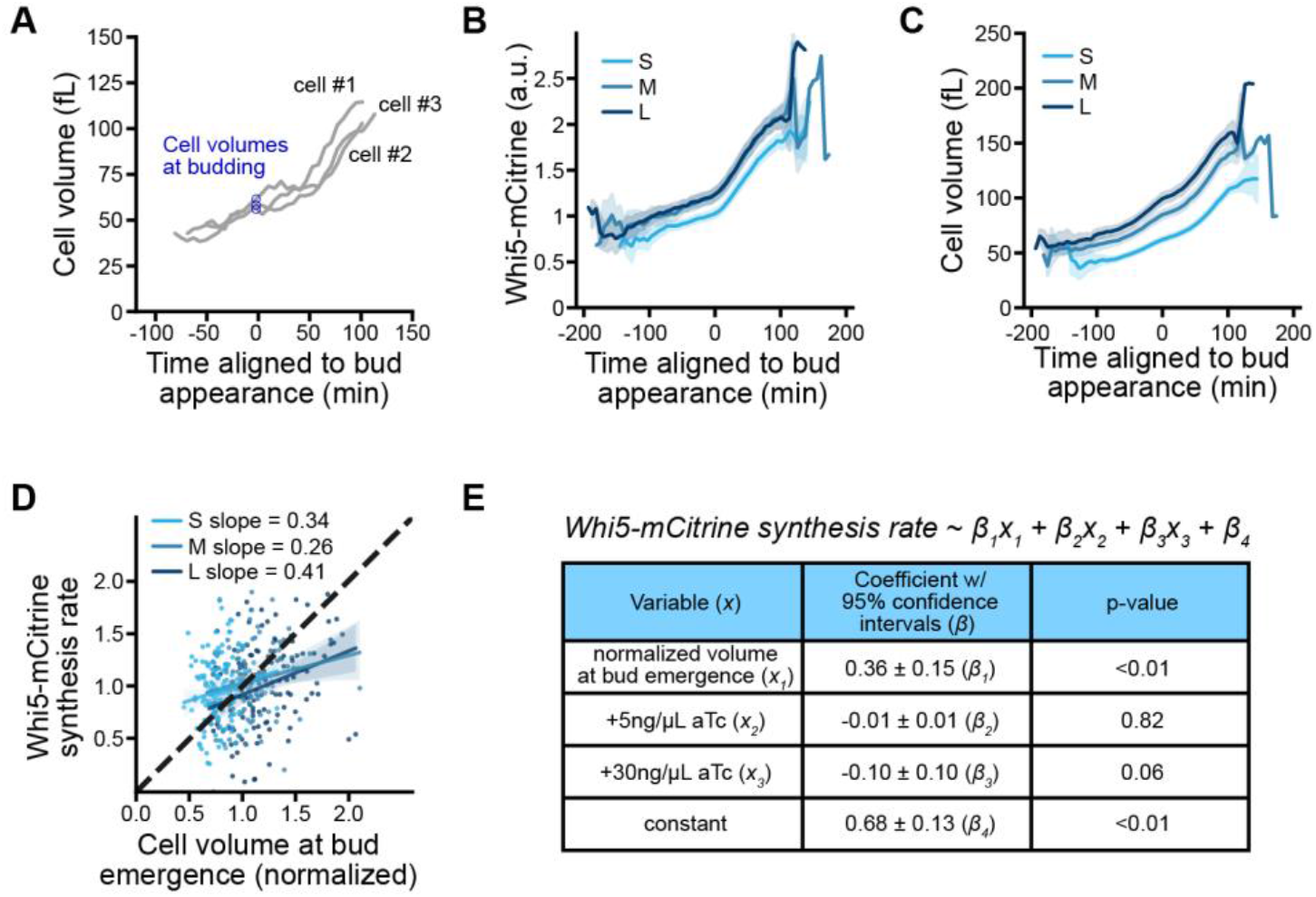
Exogenous Whi5 does not impact expression from the endogenous *WHI5* promoter. Related to Figure 1. (A) Cell volume dynamics for the corresponding cells shown in Figure 1D. (B) Average Whi5-mCitrine total fluorescence intensity dynamics aligned at budding. Small, medium, and large cells express increasing amounts of exogenous, untagged Whi5 as shown in **Figures 1F-G**. Shaded region denotes 95% confidence intervals. (small: n=140, medium: n=111, large: n=120) (C) Average cell volume dynamics, and 95% confidence intervals, for the same sets of cells shown in panel B. (D) Whi5-mCitrine synthesis rates normalized to the average rate and plotted as a function of cell size at bud emergence, which was normalized to the mean. Individual data points shown for Whi5-mCitrine synthesis rates corresponding to the binned averages shown in **Figure 1H**. The shaded regions denote the 95% confidence intervals of linear fits. (small: n=140, medium: n=111, large: n=120) (E) Multivariate linear regression statistics that did not identify a significant effect of exogenous Whi5 (medium and large cell size variables) on the synthesis rate of the endogenous Whi5-mCitrine.

**Figure S2:**
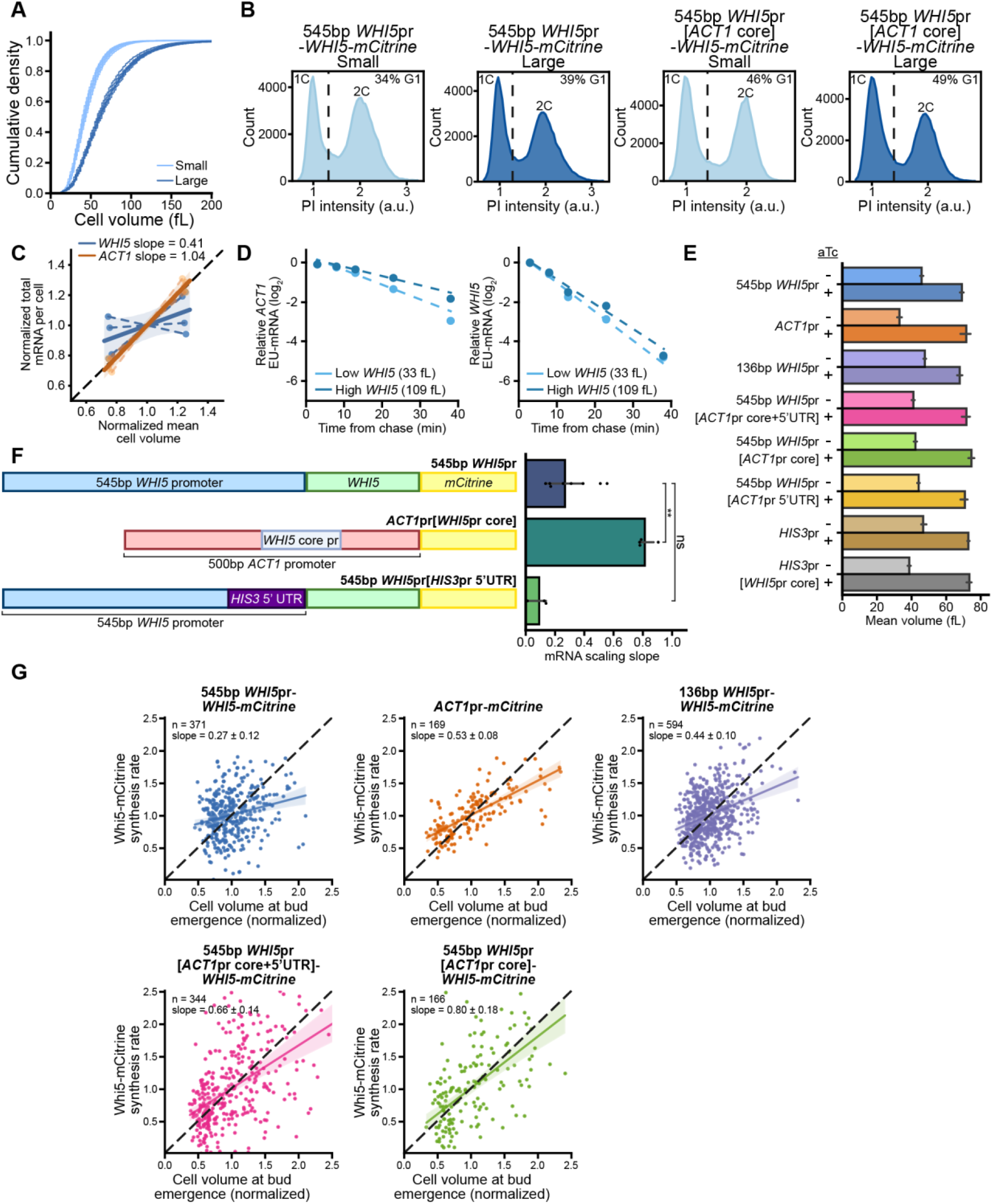
RT-qPCR analysis of cell size scaling and corroboration by single cell imaging data. Related to Figure 2. (A) Cumulative distributions of cell size measured using a Coulter counter for 9 replicates of cells exposed to 0 or 30ng/mL aTc. (B) Flow cytometry data of propidium iodide (PI) stained yeast cells for the indicated genotypes showing the distribution of DNA content used to estimate the fraction of cells in G1 phase of the cell cycle (see **methods**). (C) Spike-in normalized RNA-seq data from Swaffer et al 2023^28^ plotted to show *WHI5* and *ACT1* mRNA per cell for wild-type and larger *cln3Δ* cells. Black dashed line represents perfect size scaling where slope=1. For each gene, the mRNA values were normalized to the average of the small and large cells in each experiment. Dashed lines represent the scaling slope from each experiment, while solid lines represent a linear regression calculated from all experiments for each gene. (D) Data from Swaffer et al 2023^28^ showing the degradation of *WHI5* and *ACT1* mRNA. A spike-in normalized EU pulse-chase experiment was used to determine mRNA turnover rates in cells with low (33fL) or high (109fL) exogenous *WHI5* expression from a beta-estradiol responsive promoter. Time is aligned to EU washout (chase) following a 1-hour EU pulse. The mean (± range) of two biological replicates is shown. Dashed lines show exponential fits to the data. (E) Mean cell volumes used to calculate size scaling slopes in **Figure 2E**. Mean values were calculated from cell volume distributions measured by a Beckman Coulter Z2 Cell and Particle Counter. (545bp *WHI5*pr: n=9, *ACT1*pr: n=4, 136bp *WHI5*pr: n=8, 545bp *WHI5*pr[*ACT1*pr core+5’UTR]: n=5, 545bp *WHI5*pr[*ACT1*pr core]: n=5, 545bp *WHI5*pr[5’UTR]: n=4, *HIS3*pr: n=9, *HIS3*pr[*WHI5*pr core]: n=9) (F) Schematic representation of the promoters whose expression is examined. The core *ACT1* promoter was replaced by the core *WHI5* promoter in one strain, while the *WHI5 5’UTR* was replaced with the corresponding region of *HIS3* in another strain. Right hand side: Size scaling slope calculated as shown in **Figure 2C** for the indicated promoters. Error bars indicate the standard deviation, and each data point represents a separate biological replicate. For the indicated comparisons, ns denotes p>0.05, * p<0.05, and ** p< 0.01. (G) Whi5-mCitrine synthesis rates calculated as shown in **Figure 1D** plotted against the cell size at bud emergence. Each panel corresponds to the indicated genotype and data points correspond to single cells. The linear regression and associated 95% confidence intervals are shown. Black dashed lines represent perfect size scaling where slope = 1. Data summary is shown in **Figure 2G**. (545bp *WHI5*pr: n=371, *ACT1*pr: n=169, 136bp *WHI5*pr: n=594, 545bp *WHI5*pr[*ACT1*pr core+5’UTR]: n=344, 545bp *WHI5*pr[*ACT1*pr core]: n=166)

**Figure S3:**
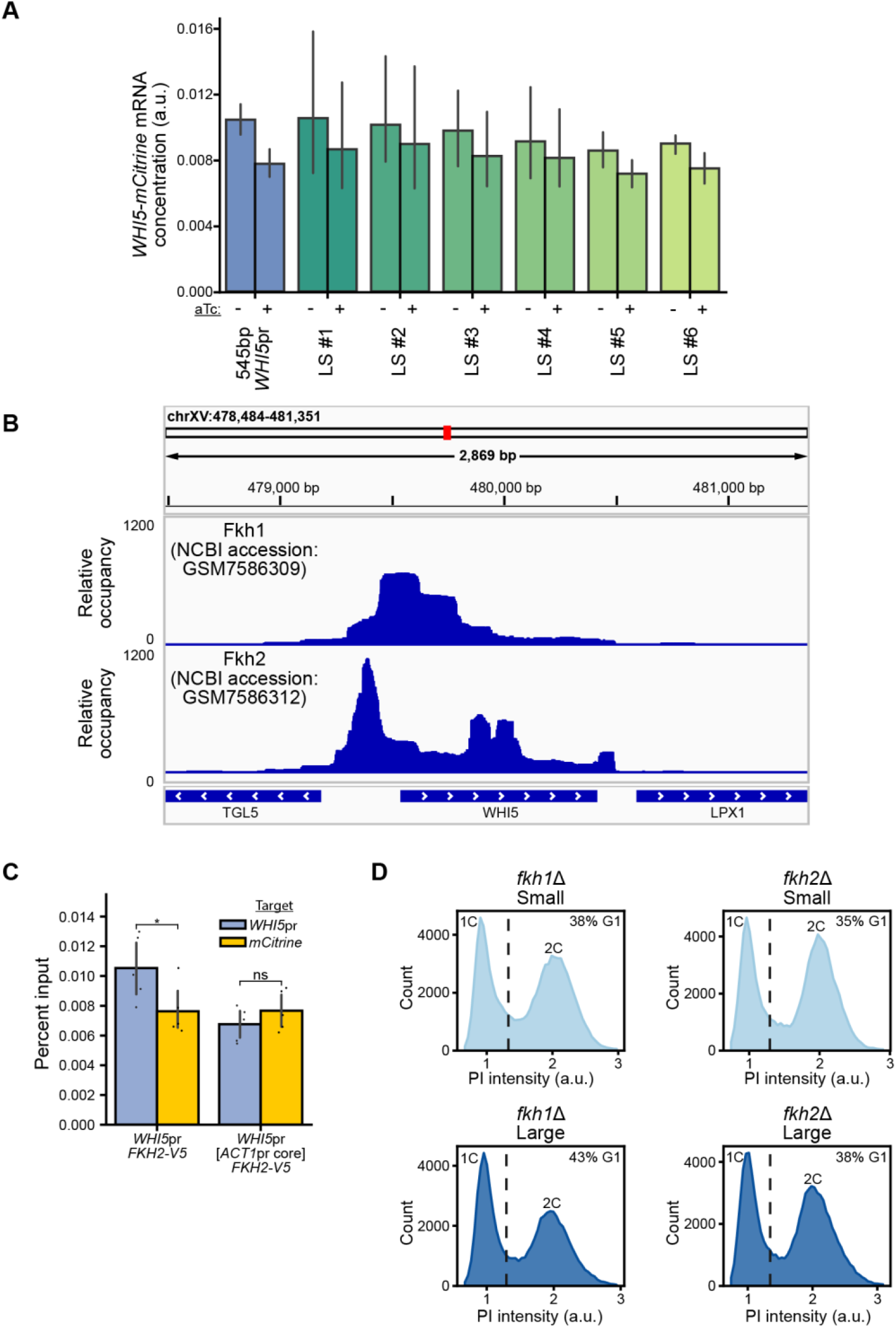
Data supporting the association of Fkh1 and Fkh2 with the *WHI5* promoter. Related to Figure 3. (A) mRNA concentrations expressed from the indicated LS mutants shown in **Figure 3A** in large and small populations of cells (±30ng/mL aTc). mRNA concentrations were measured relative to the endogenous *ACT1* mRNA. (B) Fkh1 and Fkh2 ChEC-sequencing data from Mahendrawada et al 2025^31^ showing enrichment of DNA from the *WHI5* promoter region. ChEC-seq (Chromatin Endogenous Cleavage followed by sequencing) is a genomic technique that maps where a chromatin-associated protein sits on DNA similar to the more conventional ChIP-seq technique. Bigwig files for Fkh1 and Fkh2 (GEO GSE236948) were aligned to the *S*.*cereviasiae* genome and visualized using Integrative Genomics Viewer (IGV). (C) Chromatin immunoprecipitation followed by RT-qPCR performed for the indicated strains, in which the endogenous *FKH2* gene has been fused to a V5 tag. qPCR primers target the core promoter region (see **methods**). Error bars indicate 95% confidence intervals, and each data point represents a separate biological replicate. (n=5) (D) Flow cytometry data of propidium iodide (PI) stained yeast cells for the indicated genotypes showing the distribution of DNA content used to estimate the fraction of cells in G1 phase of the cell cycle (see **methods**).

**Figure S4:**
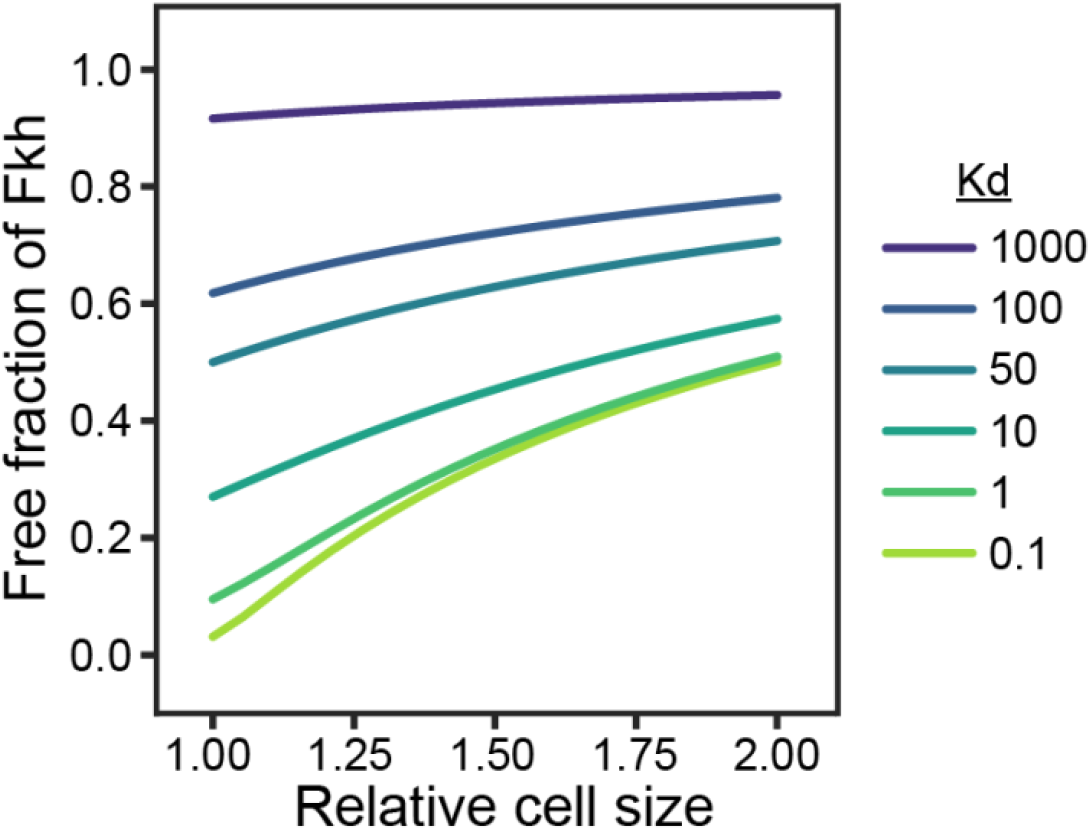
The concentration of free Fkh increases with cell size. Related to Figure 4. While the concentration of total Fkh (C_0_) remains constant, cell size increases the concentration of free Fkh because a lower proportion of the transcription factor is bound to the sites on the genome. This is due to the dilution of the fixed number of nuclear Fkh binding sites (N_s_) as the nucleus grows larger. Concentration of unbound Fkh as a function of cell volume for different values of the dissociation constant K_d_. The parameters used are C_0_=100nM and N_s_=500.

**Figure S5:**
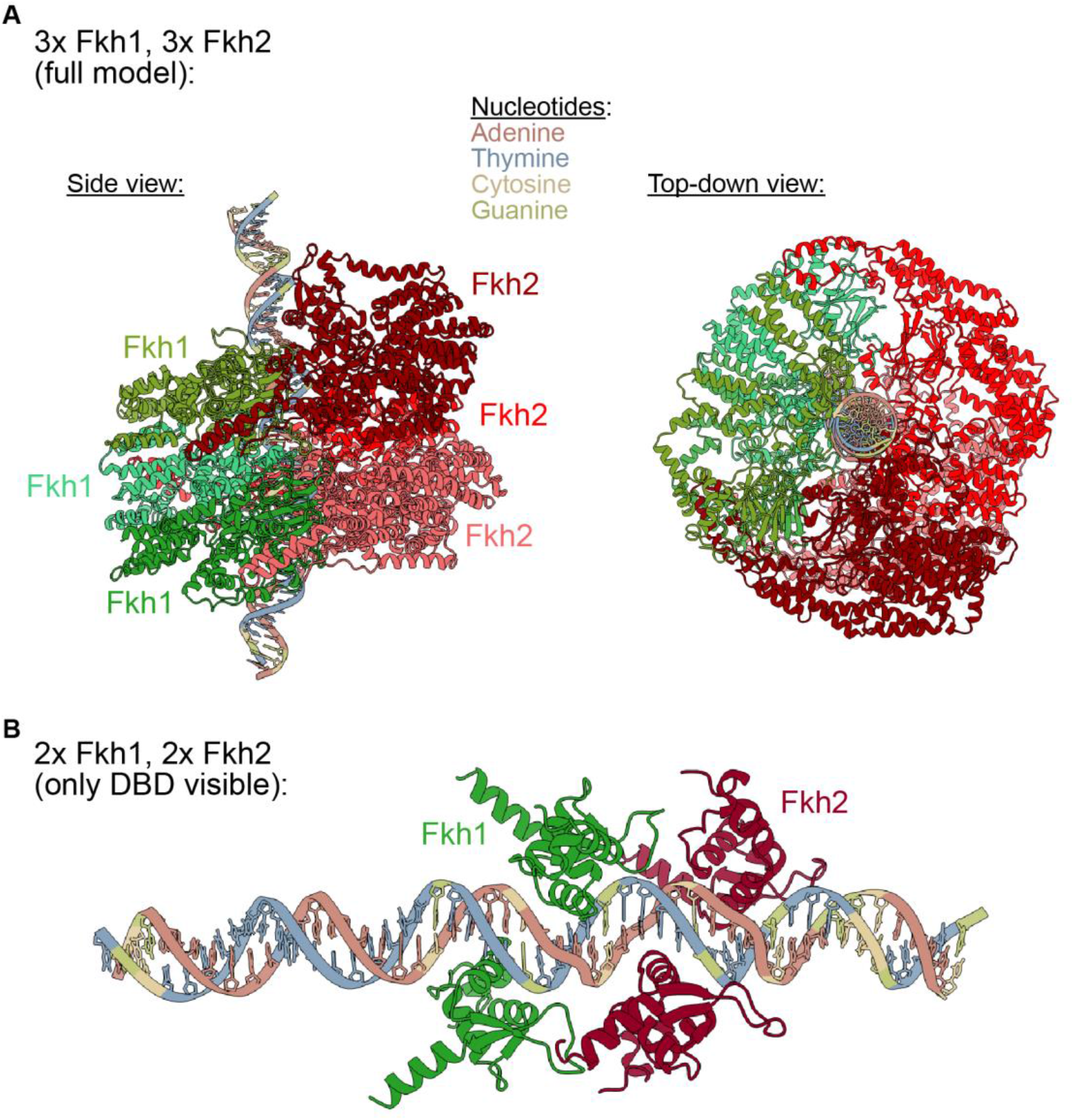
AlphaFold3 structural predictions showing Fkh1 and Fkh2 binding the core *WHI5* promoter. Related to Figure 5. (A) AlphaFold3 structural model of 3 Fkh1 proteins, 3 Fkh2 proteins, and the 51 bp *WHI5* core promoter. Fkh1 and Fkh2 molecules wrap around the core promoter, forming a multimer along each Fkh binding motif. This model displays Fkh1 and Fkh2 molecules on opposite sides of the DNA. Left: viewed laterally from DNA. Right: viewed down the double helix structure (B) DNA binding domains (DBD) of 2 Fkh1 and 2 Fkh2 proteins bound to the core *WHI5* promoter. An alternate model showing Fkh1 and Fkh2 occupying Fkh binding sites on both sides of the DNA.

**Figure S6:**
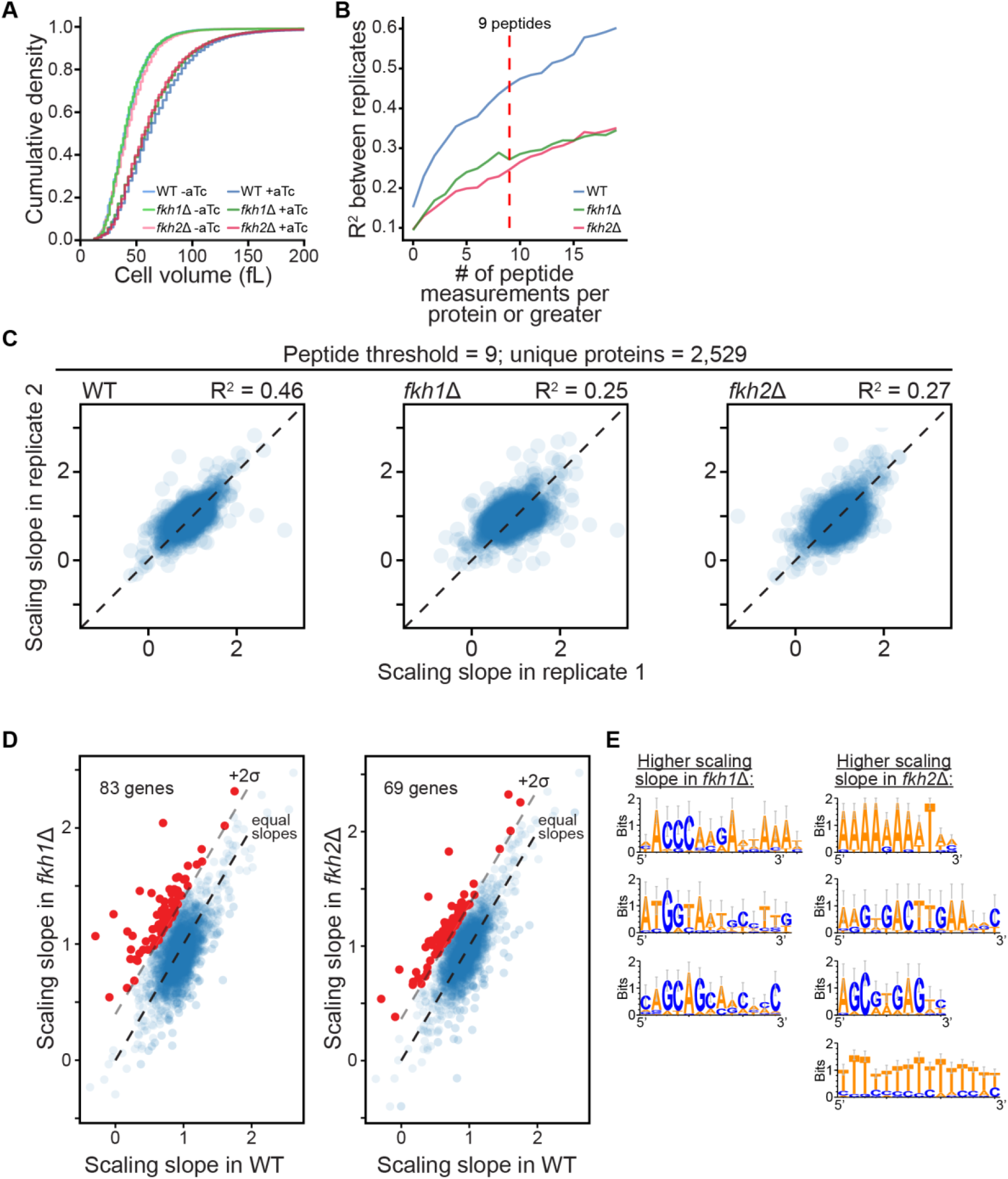
Proteomic and bioinformatic analysis of size scaling in *fkh1Δ* and *fkh2Δ* cells. Related to Figure 6. (A) Cumulative probability distributions of cell size measured using a Coulter counter for cells of the indicated genotype exposed to 0 or 30ng/mL aTc, which controls the expression of an exogenous *WHI5* allele to generate population of small and large cells, respectively. (B) The size scaling slope was measured for proteins using mass spectrometry data as described for **Figure 6B**. Slopes were compared between experimental replicates to calculate the coefficient of determination, R^2^, when only proteins for which the indicated minimum number of peptides were included. The stricter this criterion, the higher the correlation between replicate experiments, but the lower the number of proteins that were measured. To balance this tradeoff, we set this peptide threshold to 9 (red dashed line). (C) Replicate correlation of scaling slope measurements for the indicated genotypes with the peptide threshold=9. (D) Size scaling slopes calculated from protein amount estimates from the proteomics measurements for *fkh1Δ* and wild-type cells. The relative protein concentrations are taken from the mass spectrometry data and multiplied by the average cell size of the population, measured using a Coulter counter, to get the relative protein amounts. Cell size and protein amounts are normalized by the mean of the small and large cell samples for each strain. Scaling slope = (large cell protein amount - small cell protein amount)/(large cell size - small cell size). (E) Red dots indicate proteins whose slopes are more than 2 standard deviations above the trend value. (F) Sequence motifs that were enriched in the group of genes whose slopes increased upon deletion of *fkh1Δ* or *fkh2Δ* (red data points in panel D; see **methods**).

## Notes

### Competing Interest Statement

The authors have declared no competing interest.

